# Neutrophil Extracellular Trap Deposition Establishes an Immunosuppressive Environment in Non-metastatic Lymph Nodes and Facilitates Heterologous Metastatic Outgrowth

**DOI:** 10.64898/2026.10.23.761387

**Authors:** Ariane Brassard, Iqraa Dhoparee-Doomah, Shayan Hajhashemi, France Bourdeau, Betty Giannias, Jonathan Spicer, Lorenzo E. Ferri, Kim Ma, Jonathan Cools-Lartigue

## Abstract

Metastasis accounts for most cancer-related mortality, and regional lymph nodes (LNs) are among the earliest sites of dissemination for many solid tumours. Yet tumour-draining lymph nodes (TDLNs) are also sites of adaptive immune activation, raising the question of how these immune organs become permissive to metastatic growth. We investigated whether neutrophil extracellular traps (NETs), which accumulate in TDLNs before detectable tumour-cell arrival, contribute to immune conditioning of histologically tumour-negative nodes and whether the consequences of this conditioning extend beyond the initiating tumour.

Using multiplex immunofluorescence and spatial transcriptomic profiling of TDLNs from 19 patients with gastroesophageal adenocarcinoma, we found that NET positivity in non-metastatic TDLNs was associated with inferior overall survival and a distinct immunoregulatory state characterized by closer NET–Treg spatial relationships, coordinated increases in Treg-associated transcriptional programs, altered immune composition and redistribution of CD8⁺ T cells away from Tregs. NET-positive nodes also showed reduced CD8⁺ naïve/stem-like transcription, while greater peri-Treg CD8 redistribution was associated with lower cytotoxic-effector transcription. In vitro, direct NET exposure shifted human T cells toward regulatory and dysfunction-associated phenotypes and impaired tumour-cell control. NET-conditioned Tregs further suppressed CD8⁺ T-cell proliferation and granzyme B expression, while depletion of Tregs attenuated NET-associated CD8⁺ apoptosis. In tumour-bearing mice, neutrophil elastase inhibition altered activated Treg distribution across tumour-draining and contralateral LNs. Importantly, prior H59 tumour conditioning increased susceptibility to a subsequent heterologous MC38 metastatic challenge after removal of the initiating tumour; this phenotype was attenuated by TDLN resection and PAD4 deficiency. Adoptive transfer of contralateral-LN T cells from conditioned donors further identified a transferable component of this metastasis-permissive state that was sensitive to NET-directed intervention.

These findings support a model in which NET-associated nodal immune conditioning alters Treg and CD8⁺ T-cell organization and function before overt nodal metastasis. The consequences of this conditioning extend beyond the draining node and can shape host susceptibility to a subsequent heterologous tumour challenge, broadening the premetastatic-niche concept toward conditioning of the host immune state.

## 1.0 Introduction

Metastatic disease accounts for most cancer-related mortality, and regional lymph nodes (LNs) have long been recognized as among the earliest and most frequent sites of dissemination for many epithelial malignancies (1-4). Yet this tropism is not immediately intuitive: LNs are highly organized immune organs rather than intrinsically permissive sites for malignant growth (5-7). As sites of immune surveillance and adaptive immune activation, TDLNs receive tumour-derived antigens and are central to the initiation of antitumour T-cell responses (8-10). Lymphatic drainage readily explains why tumour cells reach regional LNs, but not how an organ organized to recognize peripheral antigens becomes permissive to tumour-cell survival and colonization (11, 12). Understanding the changes that occur within histologically tumour-negative TDLNs may therefore provide insight into how an immune organ becomes a site of metastatic growth and whether this conditioning has consequences beyond the node itself.

The premetastatic niche provides an established framework for understanding how tumour-derived signals alter distant tissues before malignant-cell arrival (13-15). Tumour-derived soluble factors and extracellular vesicles can remodel stromal, vascular and immune compartments, creating environments that favour subsequent metastatic establishment (14-18). Within LNs, such changes can precede detectable tumour-cell colonization and include lymphangiogenesis, vascular and stromal remodelling, altered antigen-presenting-cell function and changes in local immune composition (15, 16, 19, 20). These observations provide a compelling explanation for how an initially tumour-negative LN can become permissive to metastasis. However, the premetastatic niche is generally considered in relation to the tumour that generates it: tumour-derived signals condition a distant site from which that tumour may subsequently benefit. Whether LN conditioning can establish a broader host state that facilitates metastatic outgrowth beyond the tumour that initiated the conditioning is less clear.

The immune response to cancer develops within a host shaped by prior inflammatory experience. Tissue injury and repair involve coordinated inflammatory and regulatory programs that limit collateral damage and restore tissue homeostasis, whereas tumours can engage many of these same wound-healing processes without resolution of the initiating stimulus (21-23). Repeated or unresolved stimulation may therefore leave durable changes in immune-cell composition and function that influence the response to a subsequent challenge (24-26). Consistent with this concept, tumour-bearing hosts exhibit altered responses to challenges unrelated to the tumour itself, including infection, indicating that cancer-associated immune dysfunction can extend beyond failure of tumour rejection (27-29). We hypothesized that tumour-induced conditioning of the nodal immune environment similarly establishes a persistent immune state whose consequences are not restricted to the tumour that initiated it. Specifically, we asked whether immune cells conditioned within tumour-negative LNs could alter the response to a subsequent, antigenically distinct tumour challenge, even after removal of the initiating tumour.

Neutrophil extracellular traps (NETs) provide a plausible mechanism through which such conditioning could be established within LNs. We previously demonstrated neutrophil recruitment and NET deposition in TDLNs before detectable metastatic tumour-cell arrival (19). NETs can restrict interactions between tumour cells and cytotoxic lymphocytes, impair CD8 T-cell function through NET-associated PD-L1 and NET-DNA signalling, and promote Treg differentiation and suppressive activity through metabolic reprogramming (30-35). Preliminary observations of Treg accumulation in NET-rich regions of murine TDLNs led us to ask whether these interactions become organized into a functionally consequential immune configuration within tumour-negative nodes. We therefore examined the spatial relationships among NETs, Tregs and CD8 T cells and the consequences of NET exposure for Treg suppressive activity and CD8 T-cell function and survival. Finally, we asked whether NET-directed intervention could alter immune organization beyond the tumour-draining node and modify the consequences of prior tumour conditioning.

Here, we combined spatial profiling of human tumour-negative TDLNs, mechanistic studies of NET–T-cell interactions, and murine models of tumour conditioning and adoptive T-cell transfer. We tested whether NET-associated nodal immune conditioning alters Treg and CD8 T-cell organization and function, extends beyond the tumour-draining node, and influences the host response to a subsequent heterologous tumour challenge.

## 2.0 Materials and Methods

### 2.1 Human LN cohort and tissue microarray

Patients with histologically confirmed GEA diagnosed between April 2014 and October 2018 were identified from the McGill University Health Centre Esophageal and Gastric Data and Bio-Bank. Representative LN regions were selected following histopathological review and incorporated into a previously established LN tissue microarray. Metastatic and histologically tumour-negative LNs were classified by pathological review, with nodal metastatic status independently confirmed by two pathologists. The present study analysed metastatic and non-metastatic LN specimens from 19 patients represented on this TMA. Clinical-pathological data are summarized in **Supplemental Table 1**. The Esophageal and Gastric Data and Bio-Bank is registered under Nagano protocol MP-37-2007-856. All specimens were de-identified before analysis and participants provided written informed consent. Experimental studies were approved by the MUHC Research Ethics Board (REB 2024-10272).

### 2.2 Spatial transcriptomic and multiplex-immunofluorescence analysis

Four-micrometre TMA sections were analysed using the GeoMx Digital Spatial Profiler (NanoString). Sections were stained for CD8, FOXP3 and H3Cit, with SYTO13 used for nuclear visualization. Regions of interest were selected within each LN core for spatial transcriptomic profiling. H3Cit-positive area was quantified using HALO (Indica Labs) and used to classify LNs according to NET status. Immune-cell composition were estimated using GeoMx immune deconvolution, and differential gene and pathway analyses were performed using Partek Flow, GenePattern and gene-set enrichment analysis (36).

### 2.3 Spatial analysis of NET, Treg and CD8 organization

Canonical HALO object exports from adjacent LN sections were analysed separately for CD8/FOXP3 and H3Cit/FOXP3 staining. Because serial sections were used, cross-stain linkage was performed only at the biological LN level. Object centroids were converted to physical coordinates using the original OME-TIFF calibration in QuPath v0.6.0, and nearest-neighbour distances were calculated within each HALO Analysis Region (37).

For each CD8 cell, distance to the nearest Treg was calculated. Eligible nodes contained at least 50 CD8 cells and 10 Tregs. Among CD8 cells within 30 µm of a Treg, immediate occupancy was defined at 11–<14 µm and outer occupancy at 23–<28 µm; the CD8 clearing index was outer minus immediate occupancy. NET number, NET nearest-neighbour distance, Treg-to-nearest-NET distance, and the fraction of Tregs within 50 µm of a NET were quantified independently. GeoMx transcriptional module scores were calculated from Q3-normalized expression data as the unweighted mean of gene-level z-scores for predefined Treg- and CD8-associated gene sets; the complete gene composition and scoring procedure are provided in the **Supplementary Methods** and **Table S5**.

Node-level values were averaged within patient and NET/metastasis class. Primary non-metastatic NET+ versus NET− comparisons used an exact constrained partially paired permutation test preserving within-patient dependence; Holm correction was applied across the two prespecified CD8 endpoints. Exploratory four-state analyses compared MET−/NET−, MET−/NET+, MET+/NET+, and MET+/NET− nodes using patient-clustered models with TMA-stratified sensitivity analyses. Full computational details are provided in the **Supplementary Methods**.

### 2.4 Human blood processing and NET isolation

Peripheral blood was obtained from healthy volunteers. Neutrophils and peripheral blood mononuclear cells were isolated using established density-based methods, and T cells were enriched using negative-selection kits as appropriate (19, 38). NETs were generated from freshly isolated neutrophils by stimulation with 250 nM PMA for 4 hours at 37°C, followed by sequential centrifugation to remove cellular debris and pellet extracellular NET material. NETs were resuspended in RPMI and used immediately or stored at −80°C. Where applicable, T cells and NETs were obtained from the same donor.

### 2.5 In-vitro T-cell/NET functional assays

#### 2.5.1 T-cell NET exposure

Human T cells were cultured under CD3/CD28 stimulation with IL-2 in the presence or absence of autologous NETs for up to 5 days. T-cell subset composition and activation/exhaustion phenotypes were assessed by flow cytometry.

#### 2.5.2 Tumour-cell growth suppression assay

T cells cultured with or without NETs were subsequently cocultured with A549-GFP tumour cells at defined effector-to-target ratios. Tumour-cell growth was monitored by serial Incucyte imaging over 24 hours, with representative endpoint images obtained by EVOS microscopy.

#### 2.5.3 Treg suppression assay

CD4+CD25^high^ Tregs and CD4+/CD8+ effector T cells were isolated by flow sorting from healthy-donor PBMCs. Tregs were pre-exposed to NETs for 48 hours before coculture with CellTrace-labelled effector T cells at a 1:2 or 1:4 Treg:effector ratio under activating conditions. Effector-cell proliferation and activation were quantified by flow cytometry.

#### 2.5.4 Apoptosis assay

Bulk or Treg-depleted T-cell populations were cultured with or without NETs for 48 hours under activating conditions. Apoptosis was assessed by Annexin V-based flow cytometry to determine the relative contribution of NET exposure and Tregs to effector T-cell death.

### 2.6 Cell lines

MC38-RFP, H59-GFP and A549-GFP tumour cell lines were maintained under standard culture conditions at 37°C and 5% CO₂ in the recommended basal media supplemented with 10% fetal bovine serum and antibiotics. MC38-RFP and A549-GFP reporter lines were generated by plasmid transfection. Detailed media formulations, source information and culture conditions are provided in the **Supplementary Methods**.

### 2.7 Murine models of NET-dependent LN remodeling and metastasis

Seven- to ten-week-old C57BL/6, FOXP3-RFP, and PAD4^−/−^ mice of both sexes were used as indicated (39). All procedures were performed under institutional animal-care approval. Detailed housing conditions, reagent sources, surgical procedures, and flow-cytometry panels are provided in the **Supplementary Methods**.

#### 2.7.1 H59 flank tumour models and pharmacologic NET inhibition

Sivelestat is a selective neutrophil elastase inhibitor used here to pharmacologically limit NET formation (40, 41). C57BL/6 mice were injected subcutaneously in the left flank with 2.5×10⁵ H59-GFP tumour cells and treated daily by oral gavage with sivelestat sodium tetrahydrate (2.2 mg/kg) or an equivalent volume of vehicle.

Mice were euthanized after 20–21 days and blood, ipsilateral tumour-draining inguinal LNs and contralateral inguinal LNs were collected for flow-cytometric assessment of T-cell phenotypes. Primary tumour growth was monitored throughout treatment.

The tumour-draining LNs were also collected, fixed, and used for immunofluorescence staining to assess neutrophil (Ly6G+) recruitment, NETosis (H3Cit+), and the overall efficacy of NEi treatment.

#### 2.7.2 Induced metastasis, lymphadenectomy and PAD4-deficient model

Wild-type or PAD4^−/−^ mice first received 2.5×10^5^ H59-GFP flank tumour cells. After 21 days, the primary tumour was resected and 2.5×10^5^ MC38-RFP cells were injected intrasplenically, followed by splenectomy. A subset of wild-type mice underwent concurrent tumour-draining inguinal lymphadenectomy, while an additional control group received MC38-RFP without prior H59-GFP tumour conditioning. Mice were euthanized 14 days later for assessment of hepatic metastatic burden and immune composition.

#### 2.7.3 Adoptive transfer model

FOXP3-RFP donor mice bearing H59-GFP flank tumours were treated daily with sivelestat or vehicle. After 20 days, contralateral inguinal LNs were harvested and T cells isolated by negative selection. Recipient wild-type C57BL/6 mice received 1×10^5^ donor T cells by intrasplenic injection followed by 2.5×10^5^ MC38 cells and splenectomy; an MC38-only group served as an additional control. Recipients were euthanized 16–18 days later, and blood, LNs and liver were collected for immune profiling and assessment of hepatic metastasis.

Recipient livers were additionally processed for flow-cytometric assessment of CD4⁺ and CD8⁺ T-cell populations, FOXP3⁺ regulatory T cells, and Treg expression of PD-1 and CTLA-4. Hepatic metastatic burden was quantified from H&E-stained sections as the number of metastatic foci per unit liver-section area.

### 2.8 Flow cytometry and immunofluorescence

Single-cell suspensions from blood, LNs and other indicated tissues were analysed using BD LSR Fortessa or FACSAria instruments. Surface and intracellular staining panels were selected according to the experimental endpoint, including T-cell subset, activation/exhaustion, neutrophil and apoptosis markers. Full antibody panels and gating strategies are provided in the **Supplementary Methods**.

For immunofluorescence, formalin-fixed paraffin-embedded tissues were deparaffinized, subjected to antigen retrieval and stained with the indicated primary and fluorescent secondary antibodies. Nuclei were counterstained with DAPI, and sections were mounted in antifade medium and imaged using a Zeiss LSM780 confocal microscope.

In adoptive-transfer experiments, Tregs were identified within the CD3⁺CD4⁺ compartment by FOXP3 expression. CTLA-4 phenotype was assessed using both the proportion of CTLA-4⁺ Tregs and CTLA-4 median fluorescence intensity (MFI). Relative flow-cytometric measures were normalized within experiment as indicated.

### 2.9 Statistical analysis

Statistical analyses were performed using GraphPad Prism and R/Python where required for spatial analyses. Data are presented as indicated in the figure legends, with individual biological replicates shown whenever possible. Two-group comparisons used appropriate parametric or non-parametric tests, and comparisons involving more than two groups used one- or two-way analysis of variance with multiple-comparison procedures as appropriate.

Human spatial analyses accounted for repeated LN observations within patients. Non-metastatic NET-positive versus NET-negative comparisons used exact constrained partially paired permutation tests, with Holm adjustment across the two prespecified CD8 spatial endpoints. Exploratory four-state spatial analyses used patient-clustered models and were interpreted as hypothesis-generating. Statistical tests, sample sizes and multiplicity adjustments are specified in the corresponding figure legends and **Supplementary Methods**. A two-sided P<0.05 was considered statistically significant unless a prespecified directional hypothesis was tested.

For the pooled adoptive-transfer analysis, a Treg CTLA-4 phenotype index was calculated for each recipient as the fraction of Tregs expressing CTLA-4 multiplied by Treg CTLA-4 MFI normalized to the median PBS value from the corresponding experiment. PBS and NEi groups were compared using a two-sided exact Mann–Whitney test. Correlations between hepatic immune measures and metastatic foci density were assessed using Spearman rank correlation.

## 3.0 Results

### 3.1 NET deposition identifies an altered immunoregulatory state in tumour-negative TDLNs

We first examined the relationship between NET deposition, clinical outcome and immune organization in TDLNs from 19 patients with gastroesophageal adenocarcinoma. Metastatic (M+) and histologically non-metastatic (M−) TDLNs were analysed by multiplex immunofluorescence and GeoMx spatial profiling (Fig. 1A). NETs were detected in both M− and M+ nodes, with no significant difference in overall NET burden, whereas Tregs were more abundant in M− TDLNs (Fig. 1B).

**Figure 1.**
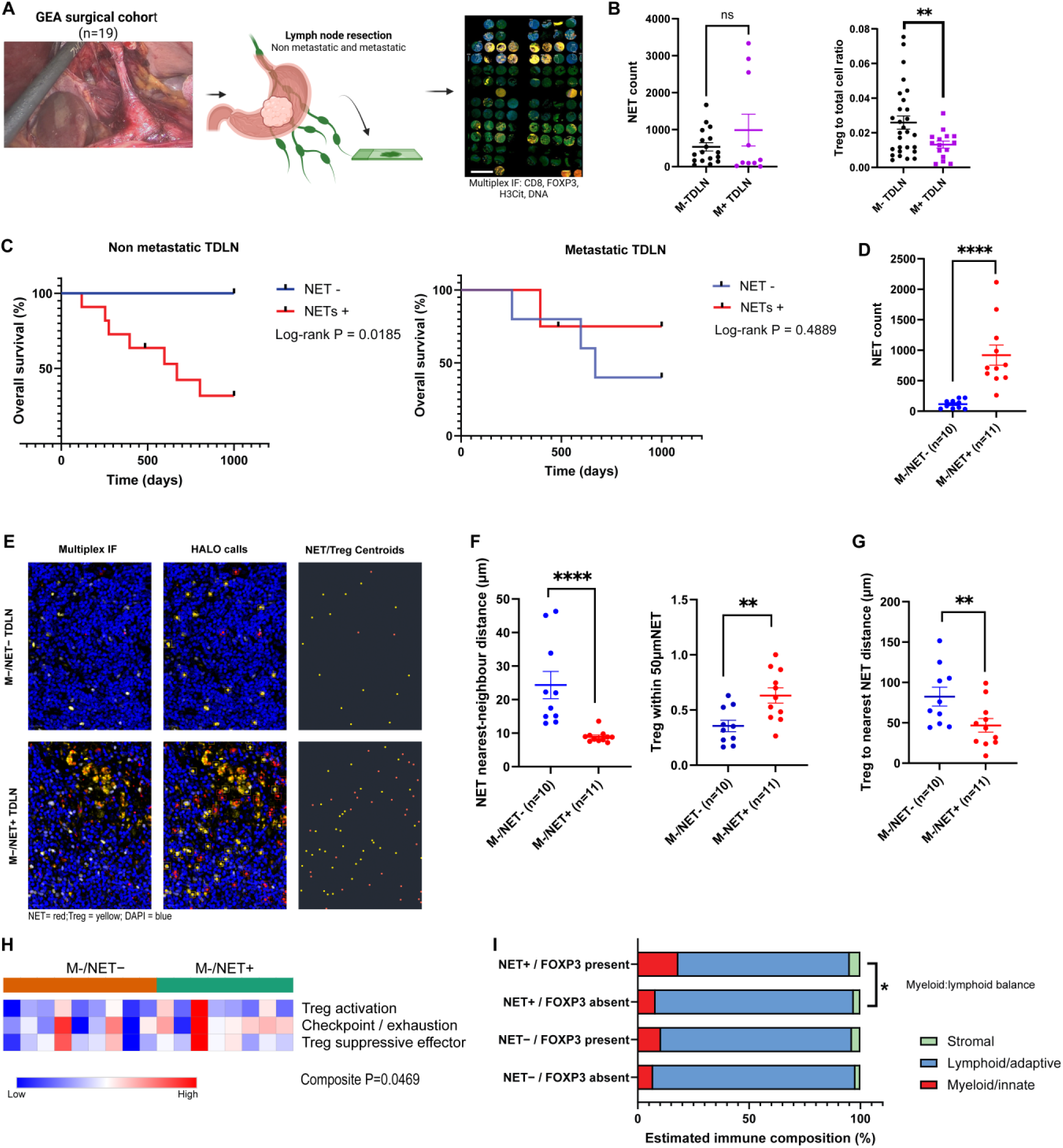
NET deposition defines an altered immune state in non-metastatic TDLNs. (A) Study workflow integrating multiplex immunofluorescence, HALO spatial analysis and GeoMx profiling of TDNLs. (B) NET burden was similar between non-metastatic (M−) and metastatic (M+) TDLNs, whereas Treg abundance was greater in M− nodes. (C) NET positivity was associated with inferior overall survival in patients with M− TDLNs (log-rank P=0.0185), but not in those with M+ TDLNs (P=0.4889). (D) M−/NET+ nodes contained greater NET burden than M−/NET− nodes. (E) Representative multiplex immunofluorescence, HALO classifications and NET/Treg centroid maps from M−/NET− and M−/NET+ TDLNs; NETs, red; Tregs, yellow; nuclei, blue. (F) M−/NET+ nodes showed shorter NET nearest-neighbour distances and a greater proportion of Tregs within 50 μm of a NET. (G) Treg-to-nearest-NET distance was shorter in M−/NET+ nodes. (H) Treg-associated transcriptional programs were coordinately increased in M−/NET+ nodes (composite P=0.0469). (I) Within M−/NET+ nodes, FOXP3-present regions showed a greater myeloid/innate-to-lymphoid/adaptive balance than FOXP3-absent regions (P=0.0278).

We next asked whether NET deposition within these nodal compartments was associated with clinical outcome. In M− TDLNs, NET positivity was associated with inferior overall survival (log-rank P=0.0185), whereas no significant association was observed in M+ TDLNs (P=0.4889; Fig. 1C). We therefore focused subsequent analyses on the immune organization of histologically tumour-negative nodes.

Within M− TDLNs, NET-positive nodes contained greater NET burden than NET-negative nodes (one-sided exact P<0.0001; Fig. 1D). Spatial analysis further showed that NETs were more closely organized in NET-positive nodes, with shorter NET nearest-neighbour distances (P<0.0001; Fig. 1E–F). Tregs were also more closely associated with NETs: a greater proportion of Tregs were located within 50 μm of a NET (P=0.0060), and median Treg-to-nearest-NET distance was shorter (P=0.0085; Fig. 1F–G).

We then examined whether this spatial relationship was accompanied by changes in Treg-associated transcription. In FOXP3-enriched regions, Treg activation, checkpoint/exhaustion-associated and suppressive-effector module scores were each directionally higher in NET-positive nodes. Although none differed significantly when considered individually (P=0.1045, P=0.1288 and P=0.1668, respectively), their prespecified equal-weight composite was increased in NET-positive TDLNs (one-sided exact P=0.0469; Fig. 1H). This coordinated increase was retained in a TMA-restricted sensitivity analysis (P=0.0357), linking greater NET–Treg proximity with a broader Treg-associated transcriptional program.

Finally, we asked whether the presence of FOXP3 was associated with differences in regional immune composition. Within NET-positive M− TDLNs, FOXP3-present regions showed greater estimated myeloid/innate representation than FOXP3-absent regions (18.4% vs 7.8%) and lower lymphoid/adaptive representation (76.7% vs 89.5%). Accordingly, the myeloid/innate-to-lymphoid/adaptive balance was higher in FOXP3-present regions (P=0.0278; Fig. 1I). The direction of this association was consistent across both TMAs, although statistical support was attenuated in the TMA-restricted sensitivity analysis.

Thus, NET deposition in histologically tumour-negative TDLNs was associated with inferior survival and identified a spatially and transcriptionally distinct immunoregulatory state characterized by NET–Treg proximity, increased Treg-associated transcription and a shift toward greater myeloid/innate representation.

### 3.2 NET-associated nodal conditioning is linked to altered CD8 spatial organization and transcriptional state

We next asked whether the NET-associated immune state in non-metastatic TDLNs extended to CD8⁺ T-cell organization and transcriptional state. Representative multiplex images and centroid maps suggested altered CD8⁺ T-cell positioning around Tregs in NET-positive compared with NET-negative nodes (**Fig. 2A**). We therefore quantified CD8⁺ T-cell distribution relative to the nearest Treg across defined peri-Treg intervals (**Fig. 2B**).

**Figure 2.**
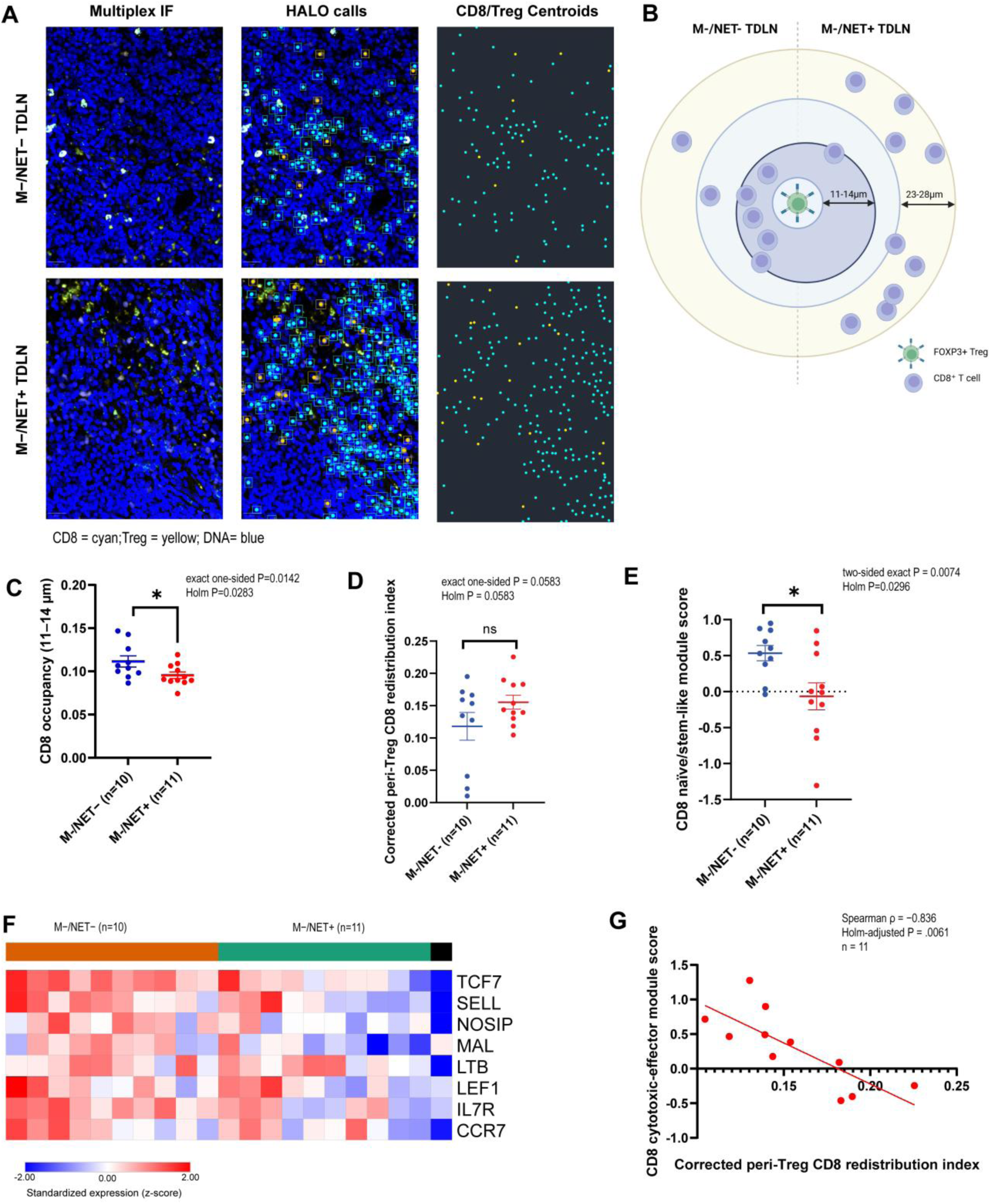
NET-positive non-metastatic TDLNs show altered peri-Treg CD8 organization and reduced stem-like CD8 transcription. (A) Representative multiplex immunofluorescence, HALO classifications and CD8/Treg centroid maps from M−/NET− and M−/NET+ TDLNs; CD8 T cells, cyan; Tregs, yellow; nuclei, blue. (B) Schematic of peri-Treg CD8 spatial analysis. (C) Immediate peri-Treg CD8 occupancy was reduced in M−/NET+ TDLNs (exact *P*=0.0142; Holm-adjusted *P*=0.0283). (D) Peri-Treg CD8 redistribution was greater in M−/NET+ nodes but did not reach statistical significance (exact *P*=0.0583; Holm-adjusted *P*=0.0583). (E) CD8 naïve/stem-like transcription was reduced in M−/NET+ TDLNs (two-sided exact *P*=0.0074; Holm-adjusted *P*=0.0296). (F) All eight genes comprising the naïve/stem-like module showed directionally lower expression in M−/NET+ nodes (directional exact *P*=0.0041). (G) Within M−/NET+ TDLNs, greater peri-Treg CD8 redistribution was associated with lower cytotoxic-effector transcription (Spearman ρ=−0.836; Holm-adjusted *P*=0.0061; *n*=11).

NET-positive nodes showed reduced CD8 occupancy within the immediate 11–<14 μm peri-Treg interval (one-sided exact *P*=0.0142; Holm-adjusted *P*=0.0283; **Fig. 2C**). Consistent with a shift away from this immediate compartment, the peri-Treg CD8 redistribution index was higher in NET-positive nodes, although the difference did not reach statistical significance (one-sided exact *P*=0.0583; Holm-adjusted *P*=0.0583; **Fig. 2D**).

This altered spatial organization was accompanied by differences in CD8-associated transcription. The naïve/stem-like module score was lower in NET-positive nodes (two-sided exact *P*=0.0074; Holm-adjusted *P*=0.0296; **Fig. 2E**), with all eight constituent genes—*TCF7, LEF1, CCR7, IL7R, SELL, MAL, LTB* and *NOSIP*—showing directionally lower expression (**Fig. 2F**; directional exact *P*=0.0041). This pattern was directionally consistent across both TMAs, although statistical support was attenuated in the TMA-restricted sensitivity analysis.

Finally, within NET-positive non-metastatic TDLNs, greater peri-Treg CD8 redistribution was strongly associated with lower cytotoxic-effector transcription (Spearman ρ=−0.836; Holm-adjusted *P*=0.0061; *n*=11; **Fig. 2G**). Thus, NET-positive tumour-negative TDLNs showed altered peri-Treg CD8 organization together with loss of naïve/stem-like transcription, while greater spatial redistribution was associated with lower cytotoxic-effector transcription.

### 3.3 NET-directed intervention alters activated Treg organization beyond the tumour-draining node

We next asked whether NET-directed intervention altered Treg organization across the nodal compartment in vivo. Tumour-bearing mice were treated with the neutrophil elastase inhibitor sivelestat (NEi) or PBS, and tumour-draining and contralateral inguinal LNs were examined (**Fig. 3A**). Immunofluorescence staining and imaging of H59-bearing mice tumour-draining LNs showed a decrease in Ly6G⁺ neutrophil recruitment in the NEi-treated groupe, as well as a reduction in H3Cit+ NETosis (**Fig. 3B**).

**Figure 3.**
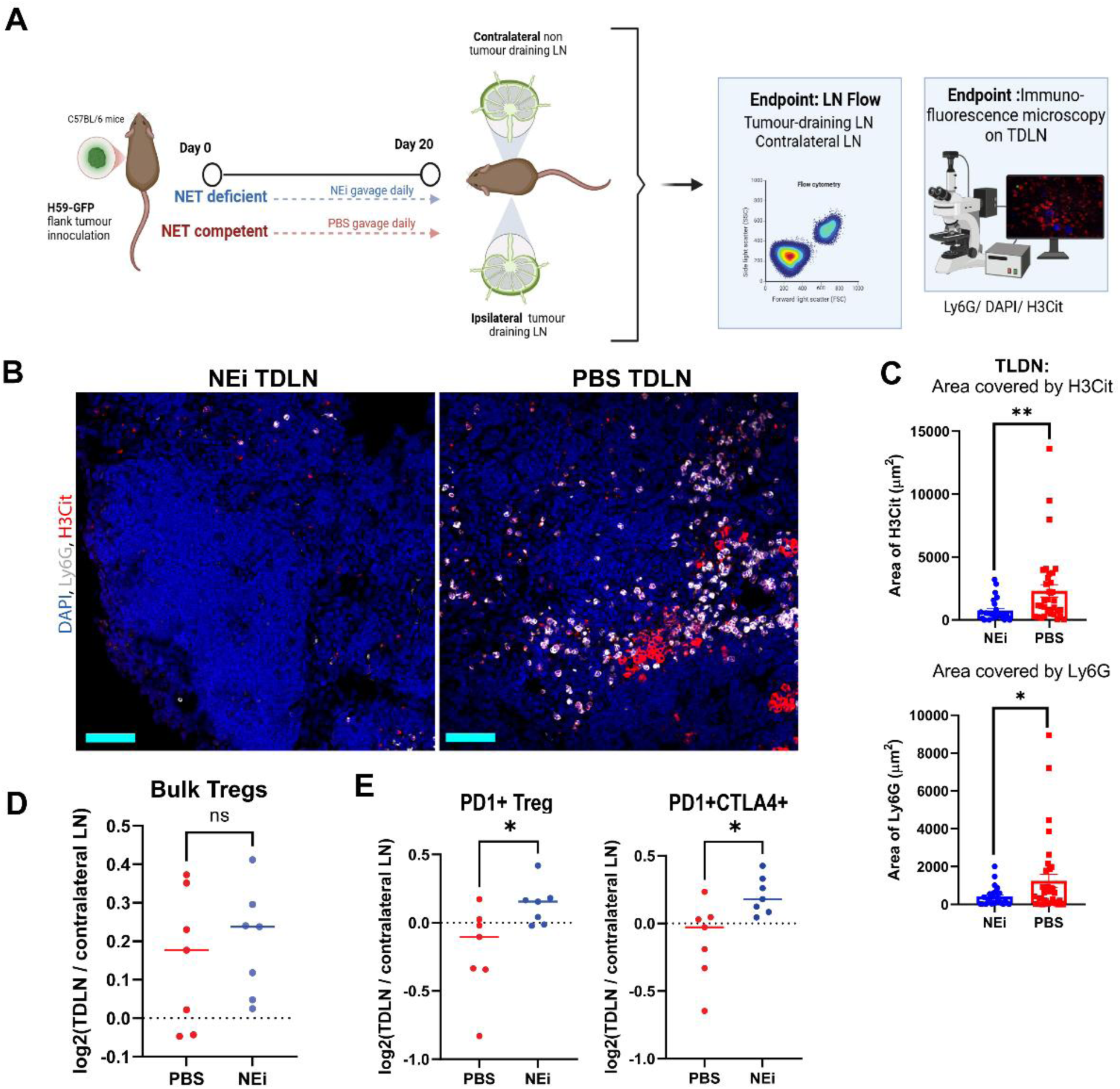
NET-directed intervention alters activated Treg organization across paired LNs. (A) Experimental design comparing tumour-draining (TDLN) and contralateral LNs in tumour-bearing mice treated with neutrophil elastase inhibitor (NEi; sivelestat) or PBS. (B) Immunofluorescence staining and imaging of the TDLN showed reduced Ly6G and H3Cit-associated signal following NE inhibition. (C) Area covered by Ly6G and H3Cit decreased in tumour-draining lymph nodes upon NEi treatment. (D) Bulk Treg distribution between paired TDLN and contralateral LNs was not significantly altered by NE inhibition. (E) NE inhibition shifted PD-1⁺ and PD-1⁺CTLA-4⁺ Tregs toward the TDLN in H59-bearing mice.

We then examined the relative distribution of Treg populations between paired tumour-draining and contralateral LNs in H59-bearing mice. NE inhibition did not significantly alter bulk Treg distribution (**Fig. 3C**). In contrast, PD-1⁺ and PD-1⁺CTLA-4⁺ Tregs shifted toward the tumour-draining LN following NE inhibition (**Fig. 3D**), indicating preferential redistribution of activated rather than bulk Tregs.

Thus, NET-directed intervention altered the distribution of activated Tregs across paired LNs in two tumour models, supporting an effect of tumour-associated NET activity on immune organization beyond the TDLN.

### 3.4 NET exposure directly shifts T cells toward regulatory/dysfunctional phenotypes and impairs tumour control

To determine whether NETs directly alter T-cell phenotype and function, peripheral-blood T cells from healthy donors were cultured for 5 days with autologous NETs, NET-preparation supernatant or control medium under activating conditions (**Fig. 4A**). NET exposure reduced the relative representation of conventional CD4⁺FOXP3⁻ and CD8⁺ T cells while increasing FOXP3⁺ Tregs (**Fig. 4B**).

**Figure 4.**
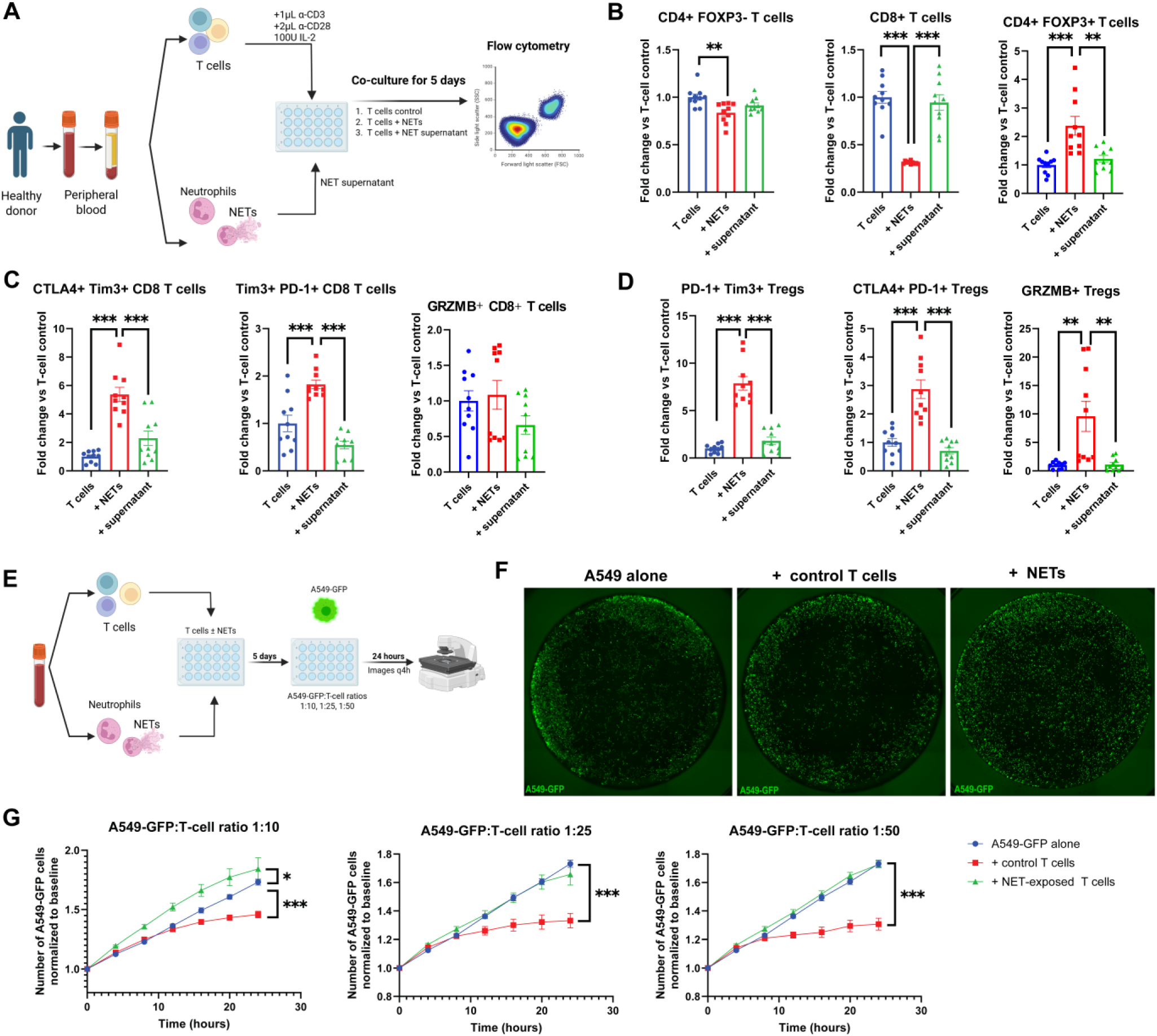
NET exposure drives dysfunctional T-cell phenotypes and impairs tumour-cell control in vitro. (A) Experimental workflow in which human peripheral-blood T cells were cultured with NETs, NET-preparation supernatant or control medium for 5 days. (B) NET exposure reduced conventional CD4⁺FOXP3⁻ and CD8⁺ T-cell representation while increasing FOXP3⁺ Tregs. (C) NET-exposed CD8⁺ T cells showed increased CTLA-4/TIM-3 and TIM-3/PD-1 co-expression. (D) NET exposure increased activated/suppressive Treg phenotypes, including PD-1⁺TIM-3⁺, CTLA-4⁺PD-1⁺ and granzyme B⁺ Tregs. (E) Experimental workflow for assessment of A549-GFP tumour-cell growth following coculture with control or NET-exposed T cells. (F) Representative A549-GFP cultures. (G) NET-exposed T cells showed impaired control of A549-GFP growth across the tested T-cell:tumour-cell ratios.

NET exposure also altered activation and dysfunction-associated phenotypes within both the CD8 and Treg compartments. Among CD8⁺ T cells, co-expression of CTLA-4/TIM-3 and TIM-3/PD-1 increased following NET exposure, whereas granzyme B expression was not increased (**Fig. 4C**). Tregs showed increased PD-1⁺TIM-3⁺ and CTLA-4⁺PD-1⁺ populations, together with greater granzyme B expression (**Fig. 4D**). These changes were largely absent in cells exposed to the NET-preparation supernatant, supporting an effect associated with the NET fraction itself.

We next asked whether these phenotypic changes translated into impaired tumour-cell control. Control or NET-exposed T cells were cocultured with A549-GFP tumour cells at defined effector-to-target ratios, and tumour-cell expansion was followed by live-cell imaging over 24 hours (**Fig. 4E–F**). Control T cells consistently suppressed A549-GFP growth, whereas NET-exposed T cells showed reduced tumour-cell control across all three tested ratios (**Fig. 4G**). At the higher tumour-cell burdens, growth in the presence of NET-exposed T cells approached that of A549-GFP cells cultured without T cells.

Thus, direct NET exposure shifted the T-cell compartment toward greater regulatory and dysfunction-associated phenotypes and reduced the capacity of T cells to control tumour-cell growth.

### 3.5 NET exposure enhances Treg-mediated suppression and contributes to CD8 T-cell loss

We next asked whether NET exposure increased the suppressive activity of Tregs. Tregs isolated from healthy-donor blood were conditioned with autologous NETs for 24 hours and subsequently cocultured with CFSE-labelled effector T cells under activating conditions (**Fig. 5A–B**).

**Figure 5.**
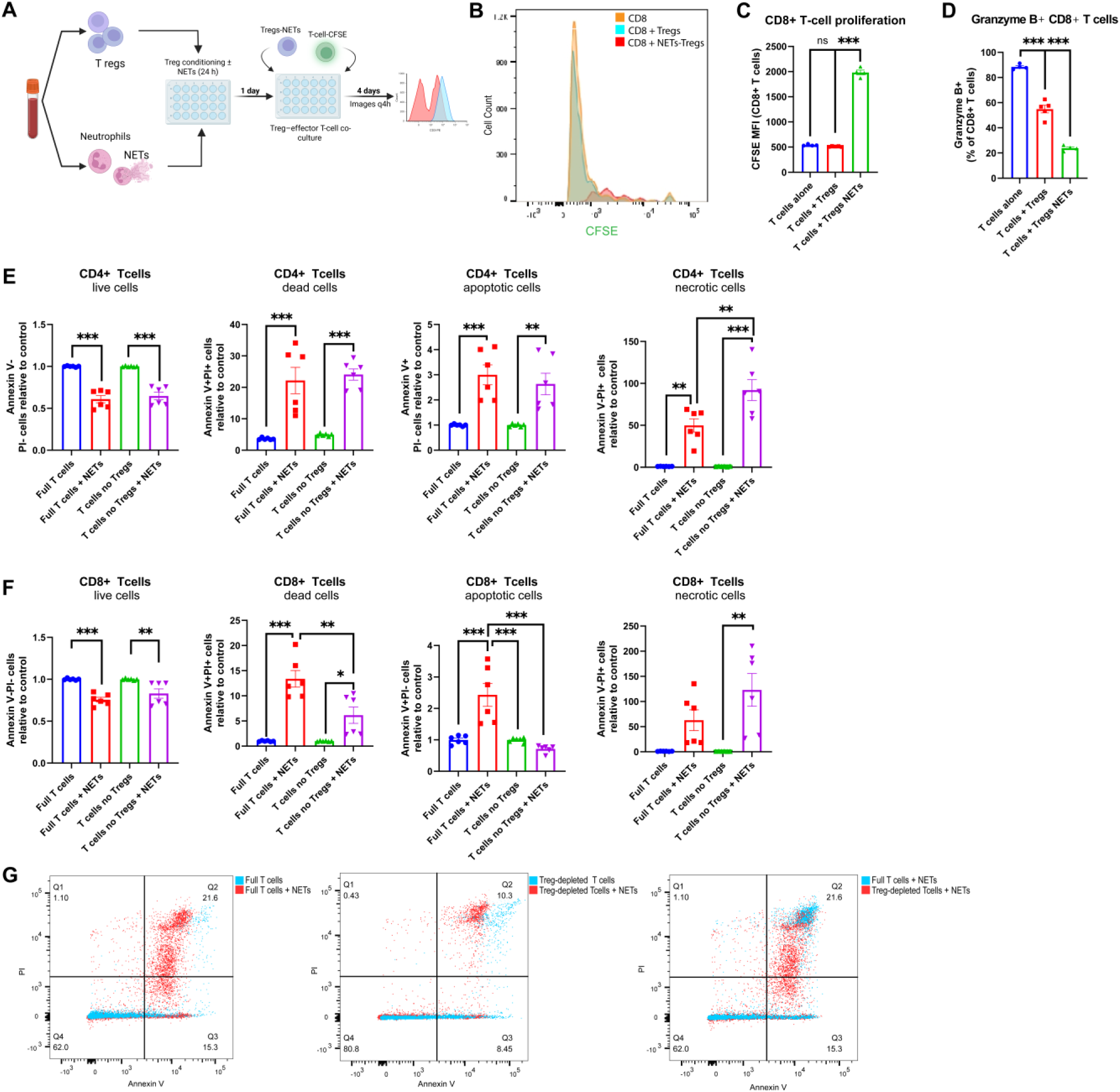
NET exposure enhances Treg-mediated suppression and contributes to CD8 T-cell apoptosis. (A) Experimental workflow in which human Tregs were conditioned with autologous NETs and subsequently cocultured with CD4⁺ and CD8⁺ effector T cells under activating conditions. (B) Representative CFSE flow-cytometry histogram demonstrating CD8⁺ T-cell proliferation following coculture with untreated or NET-conditioned Tregs. (C) CD8⁺ T cells cocultured with NET-conditioned Tregs retained greater CFSE intensity, consistent with reduced proliferation. (D) NET-conditioned Tregs reduced the proportion of granzyme B⁺ CD8⁺ T cells. (E) Annexin V/PI profiling of CD4⁺ T cells following NET exposure in complete and Treg-depleted cultures. (F) NET exposure increased CD8⁺ T-cell apoptosis in complete T-cell cultures, whereas this increase was not observed following Treg depletion. (G) Representative Annexin V/PI plots illustrating CD8⁺ T-cell death following NET exposure in complete and Treg-depleted cultures.

CD8⁺ T cells cocultured with NET-conditioned Tregs retained greater CFSE fluorescence than cells cultured alone or with untreated Tregs, consistent with reduced proliferation (*P*<0.0001; **Fig. 5C**). Granzyme B expression was also reduced in CD8⁺ T cells cocultured with NET-conditioned Tregs (*P*<0.0001; **Fig. 5D**). Thus, NET conditioning increased Treg-mediated suppression of both CD8⁺ T-cell proliferation and effector function.

We next asked whether Tregs contributed to NET-associated T-cell death. Complete T-cell populations or parallel cultures depleted of Tregs were exposed to NETs and analysed by Annexin V/propidium iodide staining. NET exposure reduced viability and increased cell-death populations among both CD4⁺ and CD8⁺ T cells (**Fig. 5E–F**). In CD8⁺ T cells, NET exposure increased the apoptotic fraction in complete T-cell cultures (*P*<0.0001), whereas no significant increase was detected after Treg depletion (*P*=0.7018; **Fig. 5F–G**).

These findings show that NET exposure enhances Treg-mediated suppression of CD8⁺ T-cell proliferation and granzyme B expression and support a contribution of Tregs to NET-associated CD8⁺ T-cell apoptosis.

### 3.6 Prior tumour conditioning establishes a TDLN- and PAD4-sensitive host state permissive to heterologous metastasis

We next tested whether tumour conditioning could establish a metastasis-permissive host state that persisted after removal of the initiating tumour. Wild-type and PAD4-deficient mice were first conditioned with subcutaneous H59-GFP tumours. After 21 days, the H59 tumour was resected and mice were challenged with the heterologous MC38-RFP tumour line by intrasplenic injection followed by splenectomy. A subgroup of wild-type mice underwent simultaneous resection of the H59 tumour-draining lymph node (TDLN), while unconditioned controls received MC38-RFP without prior H59 exposure (**Fig. 6A**). This design allowed the consequences of prior tumour conditioning to be examined independently of continued growth of the initiating tumour.

**Figure 6.**
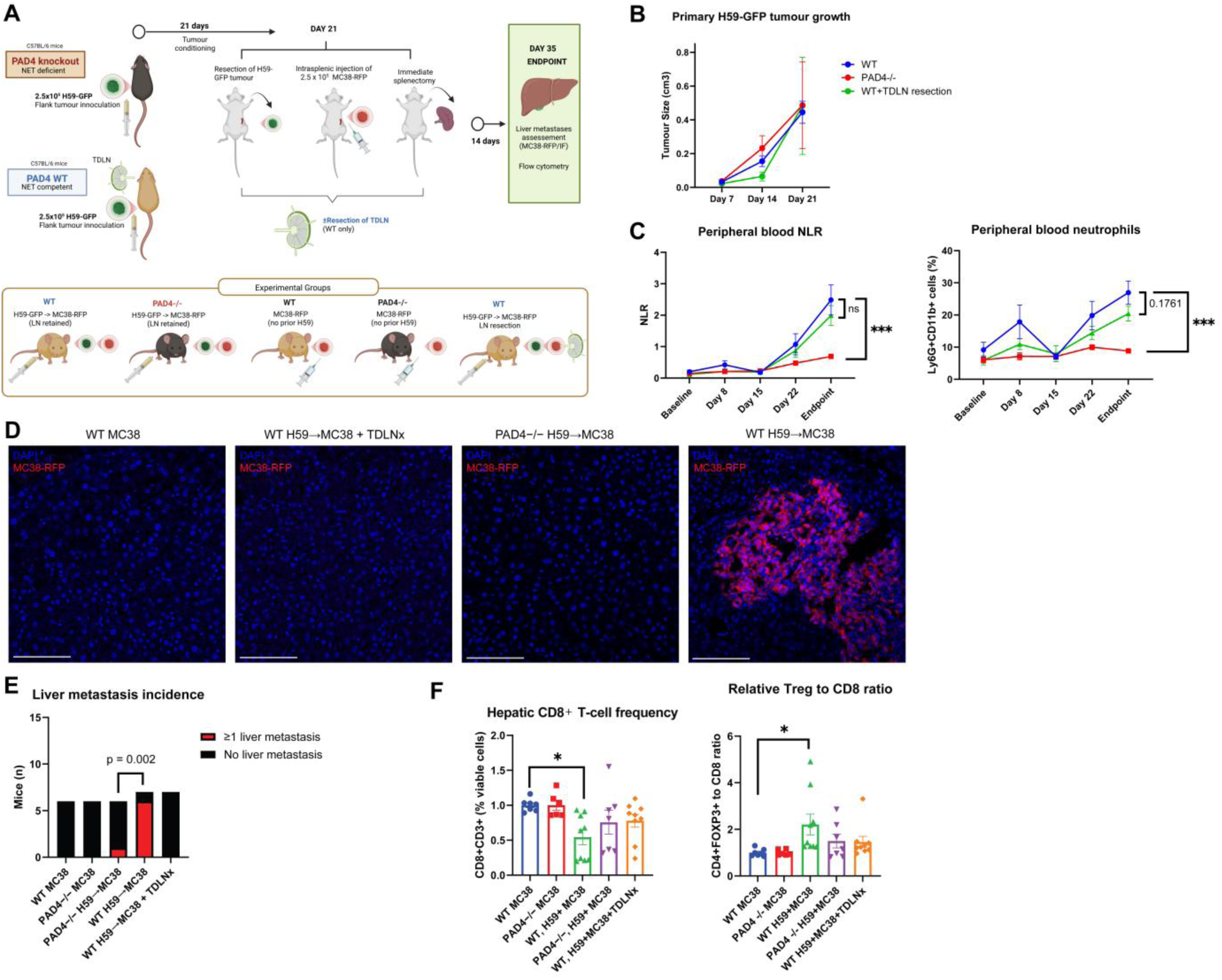
Prior tumour conditioning establishes a TDLN- and PAD4-sensitive host state permissive to heterologous metastasis. (A) Experimental model testing whether prior H59-GFP tumour conditioning alters susceptibility to a subsequent heterologous MC38-RFP metastatic challenge after removal of the initiating tumour. Wild-type and PAD4-deficient mice underwent H59 tumour resection before MC38 challenge; a subgroup of wild-type mice underwent concurrent tumour-draining lymph-node resection (TDLNx), and unconditioned controls received MC38-RFP alone. (B) Primary H59-GFP tumour growth was comparable across conditioning groups before resection. (C) H59 conditioning was accompanied by increasing peripheral neutrophilia and NLR in wild-type mice, changes that were attenuated in PAD4-deficient hosts. (D) Representative liver sections showing MC38-RFP metastatic involvement after heterologous tumour challenge. (E) Prior H59 conditioning increased hepatic metastatic incidence in wild-type mice retaining the TDLN, whereas metastasis was reduced with PAD4 deficiency, TDLN resection and in unconditioned MC38 controls (global *P*=0.0020). (F) H59-conditioned wild-type mice retaining the TDLN showed reduced hepatic CD8⁺ T-cell representation and an increased Treg-to-CD8 ratio.

Primary H59-GFP growth was comparable across the conditioning groups (**Fig. 6B**). During conditioning, wild-type mice developed increasing peripheral neutrophilia and NLR, whereas these systemic changes were attenuated in PAD4-deficient mice (**Fig. 6C**). Prior H59 conditioning markedly altered the response to the subsequent MC38 challenge. H59-conditioned wild-type mice retaining the TDLN showed the greatest hepatic metastatic incidence, whereas metastasis was reduced following TDLN resection, in PAD4-deficient hosts, and in mice challenged with MC38-RFP without prior H59 conditioning (global *P*=0.0020; **Fig. 6D–E**). Thus, enhanced susceptibility to MC38 metastasis persisted after removal of the tumour that initiated the conditioning and was attenuated by disrupting either the draining lymph node or PAD4-sensitive host processes.

The metastatic phenotype was accompanied by altered hepatic T-cell composition. H59-conditioned wild-type mice retaining the TDLN had the lowest relative CD8⁺ T-cell representation and a higher Treg-to-CD8 ratio than unconditioned MC38 controls (*P*=0.0020; **Fig. 6F**), consistent with establishment of a more immunoregulatory environment at the site of metastatic challenge.

These findings demonstrate that exposure to one tumour can condition the host to support metastatic growth of a subsequent heterologous tumour even after the initiating tumour has been removed. The attenuation of this phenotype by TDLN resection and PAD4 deficiency identifies the tumour-draining node and PAD4-sensitive processes as components of this persistent metastasis-permissive state.

### 3.7 Tumour-conditioned lymph-node T cells transfer metastatic permissiveness to naïve hosts

We next asked whether tumour-conditioned lymph-node T cells were sufficient to transmit a metastasis-permissive state to an otherwise untreated host. Contralateral LN T cells from H59-bearing donors treated with PBS or neutrophil elastase inhibitor (NEi) were transferred into naïve recipients immediately before MC38 challenge; an additional group received MC38 without T-cell transfer (Fig. 7A). Recipients of PBS-conditioned T cells developed the greatest hepatic metastatic burden, whereas metastatic foci were markedly reduced following transfer of T cells from NEi-treated donors (Fig. 7B–C).

**Figure 7.**
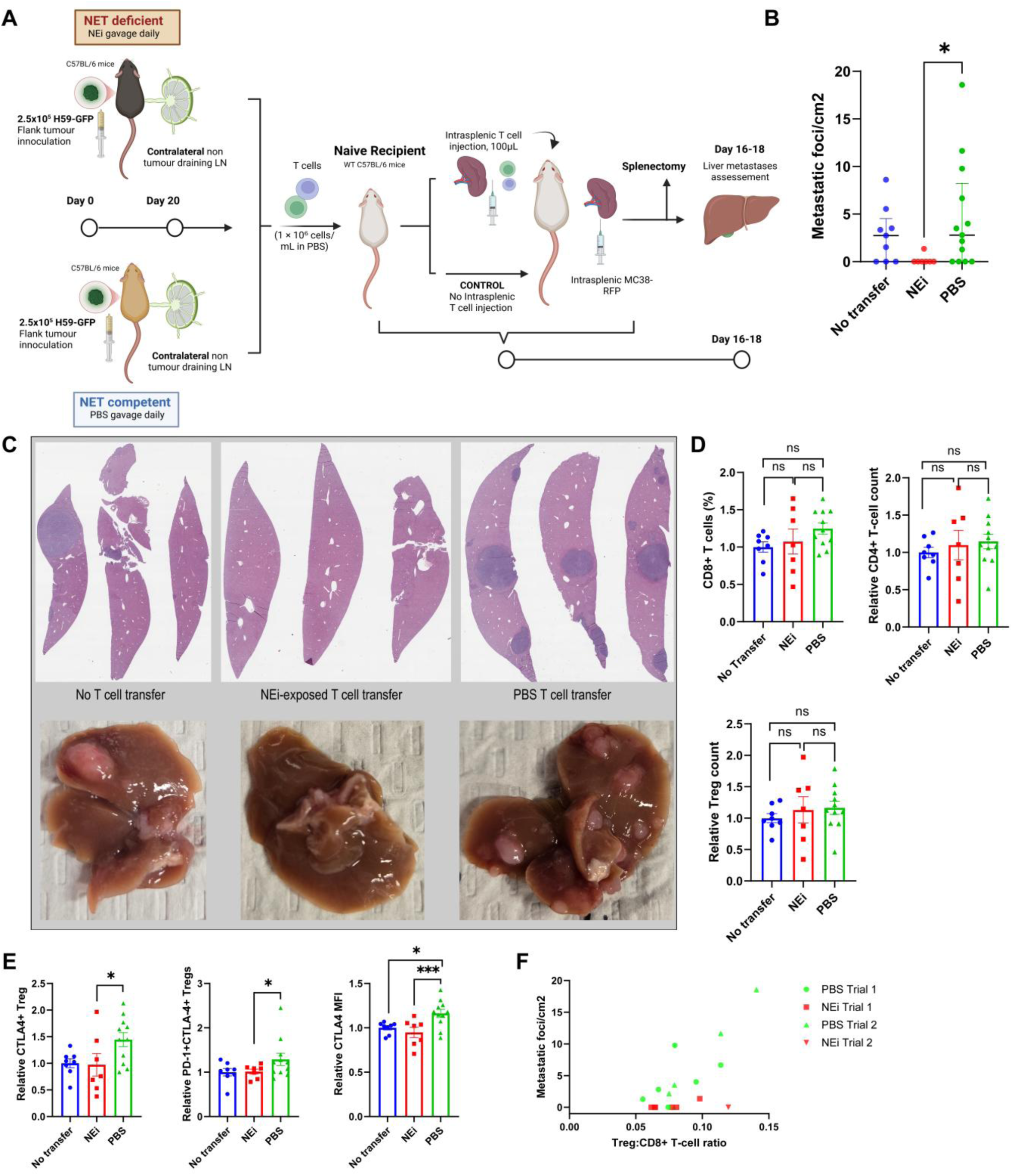
Tumour-conditioned lymph-node T cells transfer metastatic permissiveness to naïve hosts. **(A)** Adoptive-transfer model. Contralateral lymph-node T cells from H59-bearing donors treated with neutrophil elastase inhibitor (NEi; sivelestat) or PBS were transferred to naïve recipients immediately before MC38 challenge; an MC38-only group received no T-cell transfer. **(B)** Hepatic metastatic foci density in recipient mice. **(C)** Representative gross livers and H&E-stained liver sections from each recipient group. **(D)** Hepatic CD8⁺ T cells, CD4⁺ T cells and Tregs following adoptive transfer. **(E)** Hepatic CTLA-4⁺ Tregs, PD-1⁺CTLA-4⁺ Tregs and CTLA-4 expression. **(F)** Relationship between hepatic Treg:CD8⁺ T-cell ratio and metastatic foci density across two independent experiments.

This difference was not accompanied by broad changes in hepatic CD8⁺ T-cell, CD4⁺ T-cell or bulk Treg representation (Fig. 7D). Instead, it was associated with a selective shift in the regulatory phenotype: recipients of PBS-conditioned T cells showed increased CTLA-4⁺ Tregs, PD-1⁺CTLA-4⁺ Tregs and greater CTLA-4 expression compared with recipients of NEi-conditioned T cells (Fig. 7E). Across two independent experiments, variation in the hepatic Treg:CD8⁺ T-cell ratio tracked with metastatic foci density (Fig. 7F).

These findings provide direct evidence that T cells conditioned within the lymph-node environment can carry forward a metastasis-permissive immune state into a naïve recipient, and that this transferable phenotype is attenuated when NET formation is inhibited in the donor. The accompanying enrichment of CTLA-4–associated Treg features further links this transferred state to a distinct regulatory immune program rather than a nonspecific change in hepatic T-cell abundance.

## 4.0 Discussion

Our findings support a model in which tumour-associated NET formation contributes to immune conditioning of histologically tumour-negative LNs and, in doing so, alters the host response beyond the site of initial tumour drainage. In human TDLNs, NET deposition was associated with a spatially and transcriptionally distinct immunoregulatory state involving Tregs and altered CD8⁺ T-cell organization. Experimental NET exposure reproduced key elements of this phenotype, promoting regulatory and dysfunction-associated T-cell states, enhancing Treg-mediated suppression, and impairing tumour-cell control. Importantly, the consequences of tumour conditioning were not restricted to the initiating tumour: after removal of a primary H59 tumour, previously conditioned hosts showed greater susceptibility to a heterologous MC38 metastatic challenge, an effect attenuated by TDLN resection and PAD4 deficiency. The adoptive-transfer model further showed that the functional state of lymph-node T cells could modify metastatic susceptibility in recipient hosts. T cells from PBS-treated donors did not increase metastatic burden beyond MC38 challenge alone, whereas transfer of T cells from NEi-treated donors reduced metastatic outgrowth and altered the hepatic immune context. These observations suggest that TDLNs are not merely passive sites of tumour-driven remodelling, but can shape a broader host immune state in which lymph-node T-cell conditioning actively influences whether disseminated tumour cells are supported or restrained.

Regional LN involvement has traditionally been interpreted both as a marker of metastatic competence and as a potential intermediate source of systemic dissemination. Experimental lineage-tracing studies have established that tumour cells within metastatic LNs can invade nodal blood vessels, enter the circulation and seed distant organs, demonstrating that nodal metastases can provide an anatomical route for further spread (4, 42). Human phylogenetic studies, however, indicate that this is not an obligatory route: in colorectal cancer, lymphatic and distant metastases frequently arose from independent primary-tumour subclones, and a similar analysis in node-positive breast cancer identified distinct patterns of nodal and distant dissemination (43, 44). These observations suggest that the biological importance of LNs cannot be explained solely by their capacity to serve as reservoirs of disseminating tumour cells. Reticker-Flynn and colleagues provided a distinct mechanism by showing that LN colonization can induce antigen-specific Tregs and tumour-specific systemic immune tolerance that subsequently facilitates distant metastatic colonization (45). Our findings extend this framework to an earlier stage of nodal involvement, suggesting that immune conditioning is already evident in histologically tumour-negative LNs. Moreover, because prior H59 conditioning altered the response to a subsequent heterologous MC38 challenge, the consequences of this conditioning may extend beyond tolerance directed exclusively against the tumour that initiated it.

NETs appear to function not simply as extracellular traps, but as mediators between neutrophilic inflammation and adaptive immune regulation. Direct effects on lymphocytes include altered T-cell activation, physical shielding from cytotoxic lymphocytes, PD-L1-associated dysfunction and NET-DNA–mediated impairment of CD8⁺ T-cell signalling and survival (30-32, 46). NETs can also act indirectly through Tregs, promoting differentiation and suppressive activity in NASH-associated carcinogenesis and sepsis through metabolic and signalling pathways (33-35). However, this interaction is context dependent, as NETs have been reported to inhibit Treg differentiation in other inflammatory diseases (47-49). Our findings identify tumour-conditioned LNs as a setting in which the NET–Treg interaction is predominantly immunosuppressive. The attenuation of NET-associated CD8⁺ apoptosis after Treg depletion suggests that direct NET effects on CD8⁺ T cells and indirect Treg-mediated suppression likely operate in parallel rather than as a single linear pathway.

The broader implication of these findings is that metastatic susceptibility may depend not only on properties of the tumour or target organ, but also on the immune history of the host. Cancer can profoundly remodel systemic immunity across blood, marrow and peripheral tissues, including skewing haematopoiesis toward suppressive myeloid populations and altering responses to challenges unrelated to the tumour itself (27, 50, 51). Foundational studies of the premetastatic niche similarly demonstrated that primary tumours can mobilize and reprogramme bone-marrow-derived cells before tumour-cell arrival at distant sites (13, 14). These observations have largely been framed as mechanisms by which a primary tumour prepares specific tissues for its own subsequent dissemination. Our sequential tumour model raises a broader possibility: conditioning by one tumour can influence the host response to a later, biologically distinct malignant challenge after the initiating tumour has been removed. Notably, the persistence of tumour-induced immune states after resection appears to be context dependent. Systemic immune alterations can normalize rapidly following tumour removal in some models, whereas suppressive T-cell states capable of altering subsequent tumour responses have persisted after excision in others (27, 52). Our findings therefore extend the premetastatic-niche concept from preparation of a destination for an originating tumour toward conditioning of the immune host in which a subsequent malignant challenge occurs.

Antigen specificity provides one explanation for how nodal disease can influence systemic tumour immunity. Reticker-Flynn and colleagues showed that colonized LNs generate antigen-specific Tregs and tumour-specific tolerance that subsequently facilitates distant metastasis (45). More broadly, TDLNs can support either productive tumour-specific T-cell responses or tolerogenic programs, with the balance shaped by antigen presentation, regulatory-cell activity and the local immune environment (9, 53, 54). Recent work further shows that the LN itself can favour Treg-mediated suppression of tumour-specific CD8⁺ T-cell cytolytic differentiation through competition for IL-2 (ref. 55). Our models address a different question. Prior H59 conditioning altered susceptibility to a subsequent MC38 challenge after removal of the H59 tumour, while contralateral-LN T cells from conditioned donors conveyed a NET-sensitive component of this phenotype to naïve MC38 recipients. These experiments do not establish antigen independence, because T-cell clonotypes and antigen specificity were not measured. However, the heterologous challenge argues that the observed effect cannot be explained solely by tolerance directed against the initiating tumour. We therefore favour a model in which antigen recognition remains specific, while the functional context in which a new antigen is encountered has already been altered— through changes in regulatory T-cell state, effector competence and the broader immune environment

Several limitations should temper interpretation of these findings. The human analyses were performed in a small, cross-sectional cohort, and histologically tumour-negative LNs cannot be assumed to represent nodes destined to become metastatic. Spatial and transcriptional measurements were integrated at the lymph-node and patient level rather than at the same-cell level, so the data do not establish a direct three-way NET–Treg–CD8 interaction. In the murine studies, sivelestat, PAD4 deficiency and lymphadenectomy are not NET-specific perturbations and may influence additional inflammatory, immune or lymphatic processes. Finally, the sequential H59→MC38 and adoptive-transfer models establish a proof of principle for persistent and transferable tumour conditioning, but do not define its duration, antigen specificity, or the relative contributions of individual T-cell subsets.

### 4.1 Conclusion

Our findings support a model in which NET formation contributes to immune conditioning of histologically tumour-negative LNs, altering Treg and CD8⁺ T-cell organization and function before overt nodal metastasis. The consequences of this conditioning extend beyond the draining node and can influence susceptibility to a subsequent heterologous tumour challenge. Transfer of lymph-node T cells further identifies a cellular component of this conditioned state that is sensitive to NET-directed intervention. These results broaden the concept of the premetastatic niche from local preparation of metastatic sites toward conditioning of the host immune environment in which future malignant challenges occur.

## Acknowledgements

The authors acknowledge the valuable help and expertise provided by the staff of the RI-MUHC Immunophenotyping platform, the Molecular Imaging platform, the Animal Resources Division, and the Histopathology core. The authors also acknowledge the MGH foundation for their essential funding of this research.

## Authors’ contributions

AB and JCL conceptualized the project. Data acquisition for the *in vitro* experiments and murine models and corresponding analysis was performed mainly by AB, with essential aid from IDD, BG and FB. Human lymph node spatial transcriptomics analysis was done by AB, SH, JCL and KM. Manuscript writing as well as the generation of editable figures was performed by AB, JCL and KM. The manuscript was reviewed by LF and JS. All authors have read and approved the final version of the manuscript.

## Abbreviations

CFSE: Carboxyfluorescein succinimidyl ester
CTLA4: Cytotoxic T-lymphocyte associated protein 4
DAPI: 4′,6-diamidino-2-phenylindole
FOXP3: Forkhead box P3
GFP: Green fluorescent protein
GrzmB: Granzyme B
H3Cit: Citrullinated histone H3
IL: Interleukin
LN: Lymph node
MET: Metastasis
NE: Neutrophil Elastase
NEi: Neutrophil Elastase inhibitor
NET: Neutrophil Extracellular Trap
PAD4: Protein arginine deiminase 4
PBMCs: Peripheral-blood mononuclear cells
PBS: Phosphate-buffered saline
PD-1: Programmed cell death protein 1
PMA: Phorbol 12-myristate 13-acetate
RFP: Red fluorescent protein
TDLN: Tumour-draining lymph node
Tim3: T-cell immunoglobulin and mucin-domain containing-3
TMA: Tissue microarray
Treg: Regulatory T cell
WT: wildtype

