## Supplemental Methods, Tables and Figures for "Neutrophil Extracellular Trap Deposition Establishes an Immunosuppressive Environment in Non-metastatic Lymph Nodes and Facilitates Heterologous Metastatic Outgrowth"

### SUPPLEMENTARY METHODS

#### S2.2 Spatial transcriptomic and multiplex-immunofluorescence analysis

Spatial transcriptomic profiling was performed using the GeoMx Digital Spatial Profiler (NanoString) on LN tissue from the 19-patient TMA subset described in the main Methods. Four-micrometre TMA sections were stained with CD8-AF532 (clone C8/1779R; Bio-Techne), FOXP3-AF594 (clone 1054C; Bio-Techne), and an unconjugated polyclonal H3Cit antibody (Bio-Techne, NBP3-11316), detected using goat anti-rabbit IgG-Alexa Fluor 647 (Invitrogen, A-21244). SYTO 13 Green (Invitrogen, S7575) was used for nuclear visualization.

Slides were imaged on the GeoMx platform. Regions of interest (ROIs) were selected within individual LN cores with reference to the CD8 and FOXP3 fluorescence channels and corresponding tissue morphology. GeoMx segments were generated as CD8-enriched and FOXP3-enriched regions for transcriptional profiling. ROIs containing insufficient FOXP3-positive cellular content for transcriptomic analysis were classified as FOXP3/Treg-negative for the corresponding analyses. H3Cit-positive area within each core was quantified using HALO (Indica Labs), and NET status was assigned according to the prespecified H3Cit-based classification used for the study.

GeoMx analyses comprised three analytically distinct approaches: focused transcriptional module scoring, immune-cell deconvolution, and broader exploratory differential-expression/pathway analyses. These analyses were not treated as interchangeable and are described separately below.

##### S2.2.1 Focused GeoMx transcriptional module scoring

GeoMx transcriptional module scores were calculated for FOXP3-enriched and CD8-enriched segments used in **Figures 1 and 2**. The focused modules were constructed from established transcriptional markers of regulatory, activated, exhausted, naïve/stem-like and cytotoxic T-cell states reported in prior transcriptomic studies, with the exact gene composition used in the present analysis provided in **Supplementary Table S5** (refs. 1-5).

Analyses used the canonical 89-segment Q3-normalized GeoMx expression matrix. Q3-normalized target counts were transformed as $\log_2(Q3+1)$. For each gene, expression values were standardized across all 89 canonical GeoMx segments using the sample standard deviation. For each segment, the module score was calculated as the unweighted arithmetic mean of the standardized expression values of all constituent genes. All genes included in the reported modules were detected in the GeoMx dataset.

FOXP3-enriched segment scores were calculated for the following modules:

• **Treg activation:** *FOXP3, IL2RA, CTLA4, ICOS, TNFRSF18, TNFRSF4, TIGIT, IKZF2, BATF*, and *PRDM1*.
• **Checkpoint/exhaustion:** *PDCD1, LAG3, HAVCR2, TIGIT, CTLA4, TOX, ENTPD1*, and *BTLA*.
• **Treg suppressive-effector:** *FOXP3, CTLA4, IL2RA, ENTPD1, TGFB1, GZMB, TIGIT*, and *LGALS1*.

CD8-enriched segment scores were calculated for the following modules:

• **CD8 naïve/stem-like:** *TCF7, LEF1, CCR7, IL7R, SELL, MAL, LTB, and NOSIP.*
• **CD8 cytotoxic-effector:** *CD8A, CD8B, NKG7, PRF1, GNLY, GZMB, GZMK, CCL5, and CTSW.*

Where more than one GeoMx segment represented the same lymph node and segment class, replicate segment scores were averaged within lymph node. Where a patient contributed more than one lymph node within the same metastasis/NET-status category, node-level values were averaged using an unweighted arithmetic mean to obtain one patient-state value.

For the non-metastatic analyses, patient-state module scores were compared between NET-negative and NET-positive lymph nodes. The CD8 cytotoxic-effector module score was additionally correlated with the patient-level corrected peri-Treg CD8 redistribution index within non-metastatic NET-positive lymph nodes using Spearman rank correlation.

##### S2.2.2 GeoMx immune deconvolution

Immune-cell proportions were estimated using GeoMx immune deconvolution. For Figure 1I, CD8-associated regions from non-metastatic lymph nodes were stratified by NET status and FOXP3 presence/absence, generating four groups: NET-negative/FOXP3-absent, NET-negative/FOXP3-present, NET-positive/FOXP3-absent, and NET-positive/FOXP3-present.

The 18 GeoMx deconvolution cell-type estimates were grouped into three compartments. The **myeloid/innate** compartment included macrophages, mast cells, NK cells, pDCs, mDCs, classical monocytes, non-classical/intermediate monocytes, and neutrophils. The **lymphoid/adaptive** compartment included naïve B cells, memory B cells, plasma cells, naïve CD4 T cells, memory CD4 T cells, naïve CD8 T cells, memory CD8 T cells, and Tregs. The **stromal** compartment included endothelial cells and fibroblasts. Compartment percentages were calculated relative to the summed deconvolution estimates across all 18 cell types.

Where replicate GeoMx regions were available within a patient and biological state, values were averaged before patient-level comparison. The original workbook did not contain an explicit compartment-grouping formula; the grouping above was reconstructed from the 18 deconvolution columns and reproduced the Figure 1I compositions.

##### S2.2.3 Exploratory differential-expression and pathway analyses

Broader differential gene-expression and pathway analyses were performed using Partek Flow, GenePattern, and gene-set enrichment analysis (GSEA). These analyses interrogated larger externally defined pathway and cell-state signatures and were exploratory in nature.

They were analytically distinct from the focused transcriptional modules reported in **Figures 1H and 2E–G**. In particular, the focused module scores were calculated directly from prespecified compact gene sets using the standardized Q3 expression matrix, whereas the exploratory analyses evaluated larger collections of pathway and cell-state signatures with the corresponding multiple-testing framework.

Detailed coordinate-based spatial analyses of NET, Treg, and CD8 organization were performed independently from the GeoMx transcriptomic analyses and are described below.

#### **S2.3 Spatial analysis of NET, Treg and CD8 organization**

Canonical HALO object-level exports from adjacent multiplex-immunofluorescence LN sections were analysed separately. The CD8/FOXP3 section contained CD8 and FOXP3/Treg object classifications, whereas the adjacent H3Cit/FOXP3 section contained NET and FOXP3/Treg classifications. Because these markers were measured on serial sections, individual cells were not registered between sections and no same-cell NET–Treg–CD8 colocalization was inferred. Instead, sections were linked at the biological LN level using TMA, corrected patient identifier and LN designation.

Object coordinates were converted to physical distances using the original OME-TIFF calibration obtained in QuPath v0.6.0 (0.398279428 $\mu\text{m}$ per coordinate unit in X and 0.398708200 $\mu\text{m}$ in Y). The slight X/Y anisotropy was retained in Euclidean distance calculations. Nearest-neighbour analyses were performed independently within each HALO Analysis Region to prevent assignments between spatially disconnected tissue regions.

##### S2.3.1 Peri-Treg CD8 distribution

For each CD8+ T cell, the distance to its nearest Treg was calculated within the same Analysis Region. Nodes containing at least 50 CD8 cells and 10 Tregs were considered eligible. Analyses focused on CD8 cells whose nearest Treg was within 30 $\mu\text{m}$.

Within this proximal population, two prespecified Treg-centred annuli were quantified: an immediate annulus at 11–<14 $\mu\text{m}$ and an outer annulus at 23–<28 $\mu\text{m}$. Immediate occupancy was defined as the proportion of proximal CD8 cells falling within the immediate annulus, and outer occupancy as the corresponding proportion within the outer annulus. The corrected peri-Treg CD8 redistribution, or CD8 clearing index, was calculated as outer occupancy minus immediate occupancy. Positive values therefore indicate greater relative CD8 representation in the outer than immediate peri-Treg region. CD8 cells within 30 $\mu\text{m}$ but outside the two defined annuli contributed to the denominator but not to either numerator.

For descriptive visualization of the complete peri-Treg distribution, distances from 0 to <30 $\mu\text{m}$ were additionally divided into thirty 1-$\mu\text{m}$ half-open bins. The number of CD8 cells within each bin was normalized to the same <30-$\mu\text{m}$ proximal CD8 denominator. NET-positive minus NET-negative difference profiles were generated from these normalized distributions. Individual radial bins were considered descriptive and were not subjected to separate hypothesis testing.

##### S2.3.2 NET organization and NET–Treg proximity

Spatial organization within the H3Cit/FOXP3 section was evaluated independently. For each lymph node, total NET object number was recorded and the distance from each NET to its nearest neighbouring NET was calculated. The median NET nearest-neighbour distance was used as the principal summary measure of inter-NET spacing, with shorter distances indicating tighter NET spatial organization.

Treg organization was evaluated using Treg nearest-neighbour geometry, including the distance to the fifth-nearest Treg where specified in the corresponding analysis. Shorter Treg fifth-neighbour distances indicate reduced inter-Treg spacing.

For each Treg, the distance to its nearest NET was also calculated. The proportions of Tregs located within 20, 30, and 50 $\mu\text{m}$ of a NET were determined using the calibrated physical coordinates. The principal NET–Treg proximity measures used in the present analyses were the median Treg-to-nearest-NET distance and the fraction of Tregs within 50 $\mu\text{m}$ of a NET.

Because true evaluable tissue area was unavailable from the object-coordinate exports, NET density per unit tissue area was not estimated from coordinate bounding boxes. Accordingly, the reported spatial analyses address inter-object spacing and observed NET–Treg proximity rather than density-adjusted spatial attraction.

#### S2.3.3 Integration of spatial and GeoMx measurements

Spatial and GeoMx measurements were linked at the biological LN and patient level rather than at the individual-cell level. Because the multiplex-immunofluorescence measurements were obtained from adjacent sections and GeoMx measurements represented region-level transcriptional profiles, associations between spatial geometry and transcriptional programs were interpreted as cross-modality relationships within the same nodal ecology and not as evidence that the individual cells contributing to a spatial measurement were the same cells contributing to a GeoMx transcriptional signal.

For analyses relating peri-Treg CD8 organization to CD8 transcriptional state, patient-level spatial measurements were matched to the corresponding patient-level CD8-enriched GeoMx module scores within the same metastasis/NET-status category. Within non-metastatic NET-positive lymph nodes, the corrected peri-Treg CD8 redistribution index was evaluated against the CD8 cytotoxic-effector module using Spearman rank correlation.

These analyses were intended to test whether variation in LN spatial organization was associated with variation in regional transcriptional state. They do not establish causal mediation, same-cell relationships, or longitudinal progression of an individual lymph node.

#### S2.3.4 Biological unit and aggregation

Individual cells, HALO objects, GeoMx segments, and individual LN regions were not treated as independent biological replicates. Spatial metrics were first calculated at the biological LN level. Where a patient contributed more than one eligible node within the same metastasis/NET-status category, node-level measurements were averaged using an unweighted arithmetic mean.

Measurements were not weighted by total cell number, proximal CD8 denominator, tissue area, number of GeoMx ROIs, or number of HALO Analysis Regions.

The primary non-metastatic spatial analysis comprised 21 patient-state observations from 17 unique patients, including 11 NET-positive and 10 NET-negative observations. Four patients contributed observations to both NET classes, resulting in a partially paired dataset. Accordingly, NET-positive versus NET-negative comparisons used an exact dependence-preserving partially paired permutation framework rather than treating all observations as independent. Detailed permutation construction, multiplicity correction, and sensitivity analyses are described in **Supplementary Methods S2.10**.

For exploratory cross-sectional analyses, the same spatial and transcriptional measures were summarized across four prespecified nodal states: non-metastatic/NET-negative, non-metastatic/NET-positive, metastatic/NET-positive, and metastatic/NET-negative. These states represent cross-sectional LN ecologies and were not interpreted as longitudinal progression of individual lymph nodes. Full model specifications, patient-clustered inference, bootstrap procedures, and TMA-stratified sensitivity analyses are provided in **Supplementary Methods S2.10**.

### **S2.4 Human blood processing and NET isolation**

#### **S2.4.1 Neutrophil isolation and NET generation**

Peripheral-blood neutrophils were isolated from healthy volunteers as previously described. NET generation and isolation were adapted from Kaltenmeier et al (6). Briefly, $2 \times 10^6$ neutrophils were plated per well in 24-well plates and stimulated with 250 nM phorbol 12-myristate 13-acetate (PMA; Sigma-Aldrich, P8139) for 4 hours at 37°C.

Following NETosis, extracellular material was collected and centrifuged at $480 \times g$ for 5 minutes at room temperature to remove cellular debris. The resulting supernatant was centrifuged at $18,000 \times g$ for 10 minutes to pellet NET material. NETs were resuspended in 100 μL RPMI and used immediately or stored at $-80^\circ\text{C}$ for a maximum of 6 months.

#### **S2.4.2 T-cell and PBMC isolation**

T cells were isolated from peripheral blood using the EasySep Human T Cell Isolation Kit (StemCell Technologies, Cat. 17951). Where experiments required matched NET and T-cell preparations, neutrophils and T cells were obtained from the same healthy donor.

For PBMC isolation, whole blood was diluted 1:1 with PBS and layered over 15 mL lymphocyte-separation medium (LSM; Wisent, 305-010-CL). Samples were centrifuged at $800 \times g$ for 30 minutes at room temperature using minimal acceleration/deceleration to preserve the density interface. The PBMC layer was collected, washed in PBS and centrifuged at $350 \times g$ for 5 minutes before downstream use.

### **S2.5 In-vitro T-cell/NET functional assays**

#### **S2.5.1 T-cell NET exposure**

T cells and NETs were obtained from the same healthy donor where applicable. T cells were plated at $1 \times 10^6$ cells/mL in CTS OpTmizer T Cell Expansion SFM (Gibco, A1048501) in 24-well plates precoated with anti-CD3 (1 μg/mL) and anti-CD28 (2 μg/mL). Recombinant human IL-2 was added at 100 IU/mL.

Cells were cultured for 5 days in the presence or absence of NETs at 37°C and 5% CO<sub>2</sub>. T-cell subset composition and expression of activation and exhaustion markers were subsequently assessed by flow cytometry. Control conditions included untreated T cells and, where indicated, T cells cultured with the NET-preparation supernatant.

##### S2.5.2 Tumour-cell growth suppression assay

Human T cells were cultured for 5 days with or without NETs as described above. On day 4, $5 \times 10^3$ A549-GFP cells were seeded in 100 μL per well in 96-well plates and allowed to adhere overnight. On day 5, T cells were washed and added to A549-GFP cells at tumour-cell:T-cell ratios of 1:50, 1:25 and 1:10. GFP-positive tumour-cell growth was monitored using the Incucyte S3 Live-Cell Analysis System (Sartorius) at 4-hour intervals for 24 hours. Growth curves were generated from serial GFP measurements, and representative endpoint images were acquired using an EVOS M5000 Imaging System (Thermo Fisher Scientific).

##### S2.5.3 Treg suppression assay

Healthy-donor PBMCs were stained with fixable viability dye eFluor 780, CD4-Pacific Blue and CD25-APC, and CD4<sup>+</sup>CD25<sup>high</sup> cells were isolated using a BD FACSARIA Fusion cell sorter. FOXP3 enrichment within the sorted population was confirmed and these cells were used as Tregs. Tregs were cultured for 48 hours with or without NETs generated from $2 \times 10^6$ autologous neutrophils under activating conditions consisting of plate-bound anti-CD3 (1 μg/mL), plate-bound anti-CD28 (2 μg/mL), and recombinant human IL-2 (500 IU/mL).

CD25<sup>-</sup>CD4<sup>+</sup> and CD25<sup>-</sup>CD8<sup>+</sup> effector T cells were isolated by flow sorting from the same donor and labelled using a CellTrace™ CFSE Cell Proliferation Kit according to the manufacturer's instructions. Tregs and effector cells were cocultured at a 1:4 Treg:effector-cell ratio under activating conditions. Alternatively, effector T cells were isolated as described previously and not stained with the CFSE kit, and cocultured with Tregs as described above. Effector-cell proliferation and activation phenotype were assessed by flow cytometry following coculture.

##### S2.5.4 Apoptosis assay

Bulk T cells or Treg-depleted effector T-cell populations were cultured with or without NETs under activating conditions for 48 hours. Treg-depleted populations were generated by sorting CD25<sup>-</sup>CD4<sup>+</sup> and CD25<sup>-</sup>CD8<sup>+</sup> cells following staining with viability dye, CD4, CD25 and CD8. Apoptosis was assessed using an Annexin V-based Dead Cell Apoptosis Kit (Invitrogen, V13242) and quantified by flow cytometry. Parallel cultures containing or lacking Tregs were used to distinguish the effects of NET exposure alone from those occurring in the presence of Tregs.

#### S2.6 Cell lines

MC38 cells were provided by Dr. Nicole Beauchemin (McGill University) and maintained in Dulbecco's Modified Eagle Medium (DMEM; Wisent, 319-015-CL) supplemented with 10% fetal bovine serum (FBS; Wisent, 098-150) and 1% penicillin-streptomycin (Wisent, 450-201-EL). MC38-RFP cells were generated by plasmid transfection and maintained under the same conditions.

H59-GFP cells were provided by Dr. Pnina Brodt (McGill University) and maintained in RPMI-1640 medium (Wisent, 350-000-CL) supplemented with 10% FBS and 1% penicillin-streptomycin.

A549-GFP cells were generated by plasmid transfection of the parental A549 line provided by Dr. Sidong Huang (McGill University). Cells were maintained in DMEM/Ham's F-12 (1:1; Wisent, 319-075-CL) supplemented with 10% FBS and 1% penicillin-streptomycin.

All cell lines were maintained at 37°C in a humidified incubator containing 5% CO<sub>2</sub>.

### **S2.7 Murine models and tissue collection**

Seven- to ten-week-old C57BL/6 mice (Charles River Laboratories), FOXP3-RFP reporter mice (gift from Dr. Ciriaco Piccirillo, McGill University), and PAD4<sup>-/-</sup> mice (gift from Dr. Allan Tsung, University of Virginia) of both sexes were used as indicated. All procedures were performed under protocols approved by the Veterinary Authority of the Institutional Animal Care and Use Committee at the RI-MUHC.

#### **S2.7.1 H59-GFP flank tumour models and pharmacologic NET inhibition**

C57BL/6 mice were injected subcutaneously in the left flank with $2.5 \times 10^5$ H59-GFP. Mice received daily oral gavage of either sivelestat sodium tetrahydrate (2.2 mg/kg; Abcam, ab142369) or vehicle to pharmacologically limit neutrophil elastase-dependent NET formation. Vehicle-treated mice received an equivalent volume of Dulbecco's phosphate-buffered saline.

Mice were euthanized after 20–21 days. Blood, ipsilateral tumour-draining inguinal lymph nodes and contralateral inguinal lymph nodes were collected. Blood was subjected to red-blood-cell lysis using BD Pharm Lyse Lysing Buffer before flow-cytometric analysis. Lymph nodes were mechanically dissociated between glass slides, passed through cell-strainer caps and processed as single-cell suspensions for assessment of bulk and activated T-cell populations. Primary tumour growth was monitored throughout treatment.

In some cohorts, the tumour-draining lymph node was collected, fixed, and stained for Ly6G, DAPI, and H3Cit to assess neutrophil recruitment and NETosis and to compare with NEi-treated mice.

#### **S2.7.2 Induced metastasis, lymphadenectomy and PAD4-deficient model**

Wild-type and PAD4<sup>-/-</sup> C57BL/6 mice were injected subcutaneously with $2.5 \times 10^5$ H59-GFP cells in the left flank. After 21 days, the primary flank tumour was resected. During the same procedure, $2.5 \times 10^5$ MC38-RFP cells were injected intrasplenically, followed by splenectomy to prevent establishment of a splenic primary tumour.

A subgroup of wild-type mice underwent concurrent resection of the tumour-draining inguinal lymph node. An additional control cohort received MC38-RFP intrasplenic injection and splenectomy without prior H59-GFP flank-tumour conditioning.

Mice were euthanized 14 days later. Livers were collected for assessment of metastatic burden by fluorescence/confocal microscopy, and blood and tissue samples were processed for immune profiling by flow cytometry.

#### **S2.7.3 Adoptive transfer of LN T cells**

FOXP3-RFP donor mice were injected subcutaneously with $2.5 \times 10^5$ H59-GFP cells in the left flank and treated daily with either sivelestat or vehicle. After 20 days, tumour-draining and contralateral inguinal lymph nodes were harvested.

Tumour-draining lymph nodes were fixed in 10% formalin, transferred to 70% ethanol, paraffin embedded and sectioned at 4 μm for histological assessment of metastatic involvement. Contralateral lymph nodes were mechanically dissociated, and T cells were isolated using the EasySep Mouse T Cell Isolation Kit (StemCell Technologies, Cat. 19851).

Isolated donor T cells were resuspended at $1 \times 10^6$ cells/mL in PBS. Recipient wild-type C57BL/6 mice received $1 \times 10^5$ donor T cells by intrasplenic injection, followed immediately by $2.5 \times 10^5$ MC38-RFP cells. Splenectomy was then performed to prevent formation of a splenic primary tumour. An additional recipient group received MC38 cells without adoptively transferred T cells.

Recipients were euthanized 16–18 days later. Blood, inguinal lymph nodes and liver were collected. Blood and lymph nodes were processed immediately for flow cytometry, whereas liver tissue was fixed in formalin, transferred to 70% ethanol, paraffin embedded and sectioned at 4 µm for histological and immunofluorescence assessment of metastatic burden.

### S2.8 Murine sample processing and flow cytometry

Peripheral blood was subjected to red-blood-cell lysis using BD Pharm Lyse Lysing Buffer according to the manufacturer's instructions. Lymph nodes were mechanically dissociated and filtered to generate single-cell suspensions. Livers were minced into small pieces and passed through a stainless-steel mesh. The resulting pieces were suspended in RPMI and spun to pellet the larger pieces. The supernatant was collected and was resuspended, spun, and the resulting pellet was resuspended in a 37.5% Percoll gradient. The cells were spun at 1,900 rpm for 30 minutes without break. The pellet was then lysed in lysis buffer for 10 minutes and resuspended in PBS for further staining. Cells were stained with experiment-specific viability, T-cell, regulatory T-cell, neutrophil, activation and exhaustion-marker panels and acquired using a BD LSR Fortessa flow cytometer. Cell sorting, where required, was performed using a BD FACS Aria Fusion.

Full antibody panels, fluorophores, clones, dilutions and gating strategies are provided in **Supplementary Table S2**.

#### S.8.1 Tissue fixation, histology and immunofluorescence

Tissues designated for histological analysis were fixed in 10% neutral-buffered formalin for approximately 48 hours, transferred to 70% ethanol, paraffin embedded and sectioned at 4 µm.

For immunofluorescence, sections were deparaffinized, subjected to antigen retrieval and blocked for 1 hour in Tris-buffered saline containing Tween-20, 2% bovine serum albumin and 1% fetal bovine serum. Primary antibodies were applied overnight at 4°C. Following washing, fluorescent secondary antibodies and DAPI were applied for 1 hour. Slides were mounted in VectaShield antifade medium and imaged using a Zeiss LSM780 confocal microscope.

Detailed antibody information is provided in **Supplementary Table S3**.

### S2.9 Statistical Analysis

Statistical analyses were performed using GraphPad Prism and R/Python as appropriate. Individual biological replicates are shown where applicable, and summary statistics are reported as indicated in the corresponding figure legends. Two-group comparisons were performed using Welch's *t* test or non-parametric alternatives where appropriate. Experiments containing three or more independent groups were analysed using one-way ANOVA with pre-specified multiple comparisons or the corresponding non-parametric test, and experiments involving two experimental factors or repeated measurements over time were analysed using two-way ANOVA with appropriate multiple-comparison procedures. Survival distributions were evaluated using Kaplan–Meier analysis and log-rank testing. Exact tests, transformations and multiplicity corrections are specified in the corresponding figure legends. Unless otherwise stated, tests were two-sided and $P < 0.05$ was considered statistically significant.

For *in-vitro* experiments, the biological replicate was the independent healthy donor unless otherwise specified. Technical wells or repeated measurements from the same donor were not treated as independent biological replicates. Tumour-cell growth assays containing serial measurements were analysed using models accounting for treatment group and time, as specified in the figure legends.

For **murine experiments**, the individual mouse was the biological replicate. Tumour-growth experiments were analysed longitudinally using treatment and time as experimental factors. Endpoint comparisons of immune populations and metastatic burden were analysed according to the number and structure of experimental groups. No individual flow-cytometric events, image objects or metastatic lesions were treated as independent biological replicates when measurements arose from the same animal.

#### **S2.9.1 Human spatial analyses**

Primary spatial inference was restricted to non-metastatic lymph nodes and included 21 patient-stage observations from 17 unique patients, comprising 11 NET-positive and 10 NET-negative observations. Four patients contributed both a NET-positive and a NET-negative observation, whereas 13 patients contributed only one NET state. Because the dataset was therefore partially paired, group differences were evaluated using an exact constrained partially paired permutation test rather than a conventional paired or unpaired comparison.

For each of the four patients represented in both NET classes, NET-positive and NET-negative labels were either retained or exchanged within patient. Among the 13 patients contributing only one NET state, the observed allocation was preserved by assigning exactly seven observations to the NET-positive group and six to the NET-negative group in each permutation. This yielded $2^4 \times C(13,7) = 27,456$ permissible label configurations, all of which were enumerated exactly.

For each configuration, the test statistic was the difference in patient-stage group means between NET-positive and NET-negative nodes. Directional hypotheses were prespecified according to the expected spatial phenotype. For immediate peri-Treg CD8 occupancy, the alternative hypothesis was lower occupancy in NET-positive nodes; for the CD8 clearing index, the alternative hypothesis was greater clearing in NET-positive nodes. Exact one-sided P values were calculated as the proportion of permissible configurations producing a test statistic at least as extreme as the observed value in the prespecified direction. Holm correction was applied across these two prespecified CD8 spatial endpoints.

The same constrained permutation structure was applied to secondary measures of NET organization and NET-Treg proximity. Pre-specified directional hypotheses were greater NET number, shorter NET nearest-neighbour distance, a greater proportion of Tregs within 50 $\mu\text{m}$ of a NET, and shorter Treg-to-nearest-NET distance in NET-positive nodes. These secondary geometry endpoints were analysed separately and were not included in the two-endpoint Holm correction applied to the primary CD8 spatial measures.

Sensitivity analyses included a paired-only analysis restricted to the four patients contributing both NET states. All $2^4 = 16$ possible within-patient label swaps were enumerated. Because the smallest attainable one-sided P value with four pairs is $1/16 = 0.0625$, this analysis was interpreted as a directional sensitivity analysis rather than as the primary inferential test. Patient-cluster bootstrap confidence intervals were additionally generated by resampling patients with replacement while retaining together all observations contributed by each selected patient.

#### **S2.9.2 Exploratory four-state spatial analysis**

An exploratory cross-sectional analysis was performed to determine whether peri-Treg CD8 organization varied across LN states defined jointly by metastatic and NET status. Four states were specified a priori: non-metastatic/NET-negative, non-metastatic/NET-positive, metastatic/NET-positive, and metastatic/NET-negative. The first three states reflected the hypothesis that NET deposition may precede overt nodal metastasis, whereas the metastatic/NET-negative state was retained as a possible alternate or later branch. These categories were interpreted as cross-sectional states and not as longitudinal transitions of individual lymph nodes.

The analysis included 32 patient-stage observations from 18 unique patients and 41 eligible lymph nodes. The four states contained 10, 11, 4, and 7 patient-stage observations, respectively. Because individual patients could contribute lymph nodes to more than one state, all model-based inference accounted for clustering at the patient level. Immediate peri-Treg CD8 occupancy, outer CD8 occupancy, and the CD8 clearing index were evaluated using the definitions described above.

The clearing index was designated as the principal exploratory endpoint. Because the hypothesized spatial pattern predicted maximal clearing in metastatic/NET-positive nodes rather than a monotonic linear trend across all four states, a prespecified peak contrast compared the metastatic/NET-positive mean with the average of the two NET-negative endpoint states:

$$C_{\text{peak}} = \mu(\text{MET+}/\text{NET+}) - [\mu(\text{MET-}/\text{NET-}) + \mu(\text{MET+}/\text{NET-})]/2.$$

The directional alternative hypothesis was $C_{\text{peak}} > 0$. Immediate and outer CD8 occupancy were evaluated as complementary components of the same spatial phenotype. No multiplicity adjustment was applied to the single designated clearing-index peak contrast; as a sensitivity analysis, Holm adjustment was additionally calculated when immediate occupancy, outer occupancy, and clearing index were considered jointly.

Each spatial endpoint was modelled using ordinary least squares with saturated four-state indicators. Because patients could contribute multiple observations, covariance was estimated using a patient-clustered CR1 sandwich estimator. Contrast tests used 17 reference degrees of freedom, corresponding to the number of patient clusters minus one. A three-degree-of-freedom Wald F test was additionally used as an omnibus comparison of the four state means.

Uncertainty around state means and contrasts was evaluated using a patient-cluster bootstrap. Patients were sampled with replacement, and all patient-stage observations belonging to each sampled patient were retained together. Resampling continued until 200,000 valid bootstrap replicates containing all four states had been obtained. Two-sided 95% percentile confidence intervals were derived from the bootstrap distributions.

Several sensitivity analyses were performed. First, an ordinal analysis restricted to the first three states assigned scores of 0, 1, and 2 to non-metastatic/NET-negative, non-metastatic/NET-positive, and metastatic/NET-positive nodes, respectively, and tested for decreasing immediate occupancy and increasing outer occupancy and clearing index using patient-clustered models. Models were repeated with adjustment for LF-EAC-3 TMA membership.

Second, NET-positive and NET-negative nodes were compared separately within non-metastatic and metastatic strata using exact dependence-preserving randomization. Third, state-specific clearing-index values were summarized separately for LF-EAC-1 and LF-EAC-3, and an equal-TMA standardized descriptive mean and TMA-adjusted four-state model were used to assess sensitivity to TMA composition. Because several state-by-TMA cells contained few observations, these analyses were interpreted as assessments of heterogeneity rather than formal interaction testing.

Finally, a patient-fixed-effect model was used as an attenuation sensitivity analysis, and leave-one-patient-out analyses were performed by sequentially excluding each patient and recomputing the four state means and clearing-index peak contrast. Same-patient adjacent-state observations represented distinct lymph nodes and were not interpreted as longitudinal transitions of the same node. Overall, the four-state analysis was considered exploratory and hypothesis-generating rather than evidence of a defined temporal progression sequence.

### SUPPLEMENTARY TABLES

**Table S1. Table S1. Clinical and tumour characteristics of the 19-patient gastroesophageal adenocarcinoma LN cohort.**

| Characteristic | N=19 |
| --- | --- |
| Patients, n | 19 |
| Age, median (IQR)* | 66 (59.5–69.0) |
| Male sex, n/N (%) | 15/19 (78.9%) |
| NLR, median (IQR) | 3.00 (2.21–3.78) |
| NLR $\geq 4$, n/N (%) | 4/19 (21.1%) |
| pT1 | 1/19 (5.3%) |
| pT2 | 4/19 (21.1%) |
| pT3 | 13/19 (68.4%) |
| pT4 | 1/19 (5.3%) |
| pN0 | 6/19 (31.6%) |
| pN1 | 11/19 (57.9%) |
| pN3 | 2/19 (10.5%) |
| pM0 | 18/19 (94.7%); 1 missing |
| Moderate treatment response | 9/19 (47.4%) |
| No response | 4/19 (21.1%) |
| No neoadjuvant treatment | 5/19 (26.3%) |
| Response not stated | 1/19 (5.3%) |
| Overall survival, median days (IQR) | 1602 (635–2479) |
| Disease-free survival, median days (IQR) | 616 (301–1941) |
| Deaths, n/N (%) | 15/19 (78.9%) |

**Table S2. Flow-cytometry antibody panels across murine and human experiments**

| **Murine lymph node panel** |  |  |  |  |
| --- | --- | --- | --- | --- |
| **Target** | **Fluorophore** | **Clone** | **Concentration** | **Intracellular** |
| Fixable Viability Dye | eFl780 | --- | 1:1000 | No |
| CD3 | PE-Cy-7 | 145-2C11 | 1:100 | No |
| CD4 | eFluor450 | RM4-5 | 1:200 | No |
| CD8 | BV650 | 53-6.7 | 1:200 | No |
| PD-1 | BV605 | 29F.1A12 | 1:100 | No |
| PD-1 | FITC | MIH4 | 1:100 | No |
| PD-1 | FITC | 29F.1A12 | 1:100 | No |
| Tim3 | PE | F38-2E2 | 1:100 | No |
| CTLA4 | BUV737 | UC10-4B9 | 1:100 | Yes |
| CTLA4 | APC | UC10-4B9 | 1:100 | Yes |
| Granzyme B | BV510 | GB11 | 1:100 | Yes |
| FOXP3 | PE-eFl610 | FJK-16s | 1:100 | Yes |
| **Murine blood panel** |  |  |  |  |
| Fixable Viability Dye | eFl780 | --- | 1:1000 | No |
| CD3 | PE-Cy-7 | 145-2C11 | 1:100 | No |
| CD4 | eFluor450 | RM4-5 | 1:200 | No |
| CD8 | BV650 | 53-6.7 | 1:200 | No |
| Ly6G | AF647 | 1A8 | 1:300 | No |
| CD11b | BV510 | M1/7 | 1:300 | No |
| FOXP3 | PE-eFl610 | FJK-16s | 1:200 | Yes |
| **Human blood panel** |  |  |  |  |
| Fixable Viability Dye | eFl780 | --- | 1:1000 | No |
| CD3 | BV650 | UCHT1 | 1:100 | No |
| CD4 | PB | RPA-T4 | 1:200 | No |
| CD8 | AF700 | RPA-T8 | 1:200 | No |
| PD-1 | FITC | MIH4 | 1:100 | No |
| PD-1 | FITC | 29F.1A12 | 1:100 | No |
| Tim3 | PE | F38-2E2 | 1:100 | No |
| CTLA4 | BUV737 | UC10-4B9 | 1:100 | Yes |
| CTLA4 | APC | UC10-4B9 | 1:100 | Yes |
| Granzyme B | BV510 | GB11 | 1:100 | Yes |
| FOXP3 | PE-eFl610 | FJK-16s | 1:100 | Yes |

**Table S3. Antibodies and staining conditions used for immunofluorescence microscopy**

| Target | Species | Colour | Clone | Catalogue number | 1° or 2° antibody | Concentration |
| --- | --- | --- | --- | --- | --- | --- |
| **Ly6G** | Rat | AF647 | 1A8 | 127610 | 1° | 5µg |
| **MPO** | Goat | Unconjugated | Polyclonal | AF3667 | 1° | 5µg |
| **H3Cit** | Rabbit | Unconjugated | Polyclonal | ab5103 | 1° | 5µg |
| **H3Cit** | Rabbit | Unconjugated | Polyclonal | NBP3-11316 | 1° | 1 to 100 |
| **CD8** | Rabbit | AF532 | C8/1779R | NBP2-54595AF532 | 1° | 1 to 50 |
| **FOXP3** | Rabbit | AF594 | 1054C | IC8214T-100UG | 1° | 1 to 20 |
| **Anti-rabbit IgG** | Goat | AF647 | Polyclonal | A-21244 | 2° | 1 to 400 |
| **Nuclear dye** | -- | SYTO™ 13 Green dye | -- | S7575 | -- | 500nM |
| **Nuclear dye** | -- | DAPI | -- | D21490 | -- | 1 to 2000 |

**Table S4. GeoMx-derived transcriptional and immune-composition measures used in Figures 1 and 2**

| **Figure panel** | **GeoMx measure** | **Segment / AOI** | **Biological interpretation** | **Calculation** | **Aggregation / inferential unit** |
| --- | --- | --- | --- | --- | --- |
| **Fig. 1H** | Treg activation module | FOXP3-enriched | Activated/engaged Treg transcriptional state | Q3-normalized counts were transformed as $\log_2(Q3+1)$. Each gene was standardized across all 89 GeoMx segments, and the module score was calculated as the unweighted mean of the constituent gene z-scores. | Replicate segments were averaged within node and then within patient and ecological class; patient was the inferential unit. |
| **Fig. 1H** | Checkpoint/exhaustion module | FOXP3-enriched | Inhibitory/checkpoint-associated T-cell transcriptional state | Q3-normalized counts were transformed as $\log_2(Q3+1)$. Each gene was standardized across all 89 GeoMx segments, and the module score was calculated as the unweighted mean of the constituent gene z-scores. | Replicate segments were averaged within node and then within patient and ecological class; patient was the inferential unit. |
| **Fig. 1H** | Treg suppressive-effector module | FOXP3-enriched | Regulatory/suppressive effector transcriptional program | Q3-normalized counts were transformed as $\log_2(Q3+1)$. Each gene was standardized across all 89 GeoMx segments, and the module score was calculated as the unweighted mean of the constituent gene z-scores. | Replicate segments were averaged within node and then within patient and ecological class; patient was the inferential unit. |
| **Fig. 1I** | Immune-cell composition | CD8-associated regions stratified by NET status and FOXP3 presence/absence | Relative immune/stromal composition of the sampled GeoMx regions | GeoMx immune deconvolution estimates were summarized into **myeloid/innate, lymphoid/adaptive and stromal** compartments. | Displayed by NET-/NET+ × FOXP3 absent/present state. Individual GeoMx deconvolution cell types were collapsed into |
|  |  |  |  |  | myeloid/innate, lymphoid/adaptive and stromal compartments as defined in Supplementary Table S6 |
| **Fig. 2E–F** | CD8 naïve/stem-like module | CD8-enriched | Naïve/stem-like or less differentiated CD8 transcriptional state | Q3-normalized counts were transformed as $\log_2(Q3+1)$. Each gene was standardized across all 89 GeoMx segments, and the module score was calculated as the unweighted mean of the constituent gene z-scores. | Replicate segments were averaged within node and then within patient and ecological class; patient was the inferential unit. |
| **Fig. 2G** | CD8 cytotoxic-effector module | CD8-enriched | CD8 cytolytic/effector transcriptional machinery | Q3-normalized counts were transformed as $\log_2(Q3+1)$. Each gene was standardized across all 89 GeoMx segments, and the module score was calculated as the unweighted mean of the constituent gene z-scores. | Replicate segments were averaged within node and then within patient and ecological class; patient was the inferential unit. Figure 2G correlates this score with the corrected peri-Treg CD8 redistribution index in M<sup>−</sup>/NET<sup>+</sup> TDLNs. |

**Table S5. Gene composition of GeoMx transcriptional modules**

| Figure panel | Module | Genes | n |
| --- | --- | --- | --- |
| Fig. 1H | Treg activation | *FOXP3, IL2RA, CTLA4, ICOS, TNFRSF18, TNFRSF4, TIGIT, IKZF2, BATF, PRDM1* | 10 |
| Fig. 1H | Checkpoint / exhaustion | *PDCD1, LAG3, HAVCR2, TIGIT, CTLA4, TOX, ENTPD1, BTLA* | 8 |
| Fig. 1H | Treg suppressive effector | *FOXP3, CTLA4, IL2RA, ENTPD1, TGFB1, GZMB, TIGIT, LGALS1* | 8 |
| Fig. 2E–F | CD8 naïve / stem-like | *TCF7, LEF1, CCR7, IL7R, SELL, MAL, LTB, NOSIP* | 8 |
| Fig. 2G | CD8 cytotoxic effector | *CD8A, CD8B, NKG7, PRF1, GNLY, GZMB, GZMK, CCL5, CTSW* | 9 |

**Table S6. GeoMx immune deconvolution cell types grouped into collapsed immune and stromal categories for Figure 1I**

| Collapsed category | GeoMx deconvolution cell types |
| --- | --- |
| Myeloid/innate | macrophages; mast; NK; pDCs; mDCs; monocytes.C; monocytes.NC.I; neutrophils |
| Lymphoid/adaptive | B.naive; B.memory; plasma; T.CD4.naive; T.CD4.memory; T.CD8.naive; T.CD8.memory; Treg |
| Stromal | endothelial.cells; fibroblasts |

### SUPPLEMENTARY FIGURES

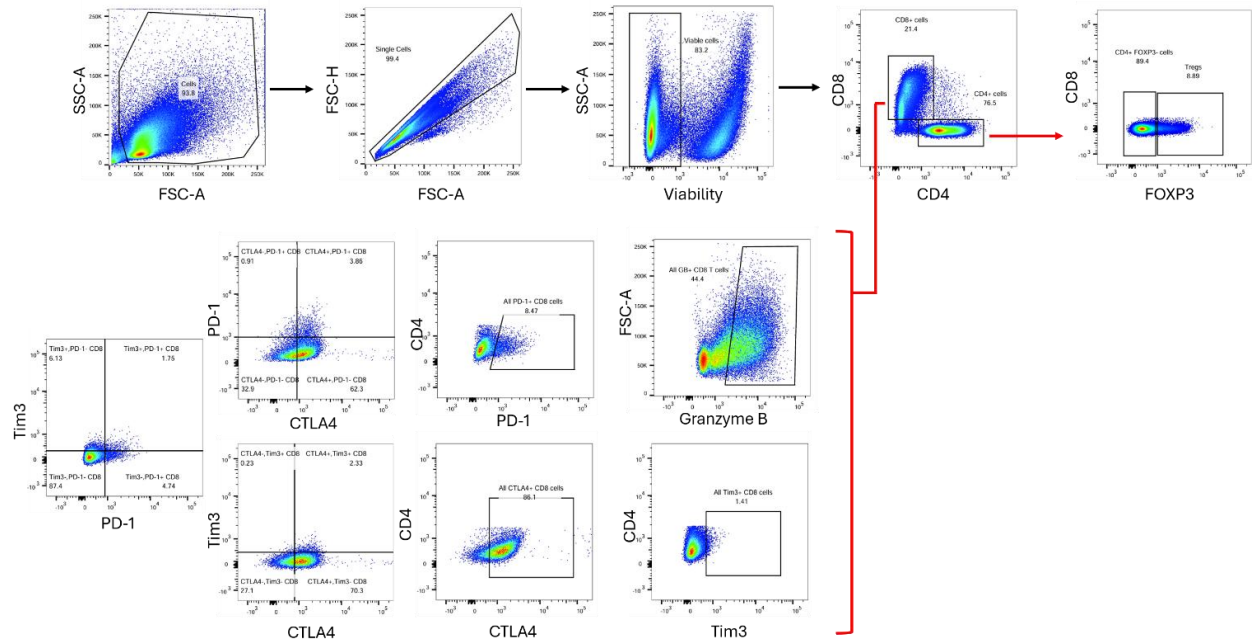

**Figure S1. Flow cytometry gating strategy for the human T cell exhaustion panel.** The gating strategy of the PD-1, CTLA4, Tim3, and granzyme B expression in the CD8<sup>+</sup> T cell population was also applied identically to the regulatory T cells and CD4<sup>+</sup> T cell population.

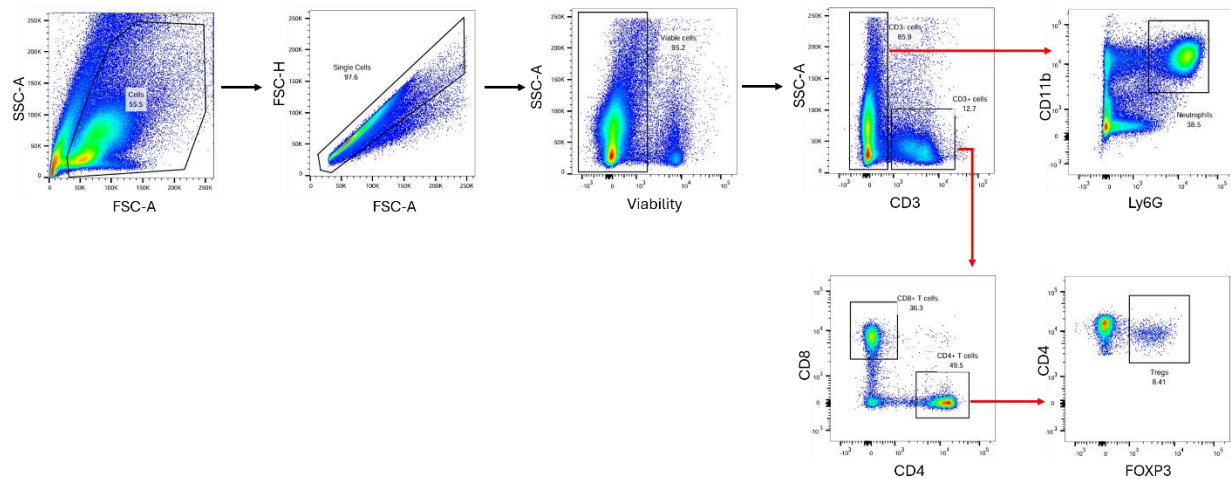

**Figure S2. Flow cytometry gating strategy for the murine blood panel.**

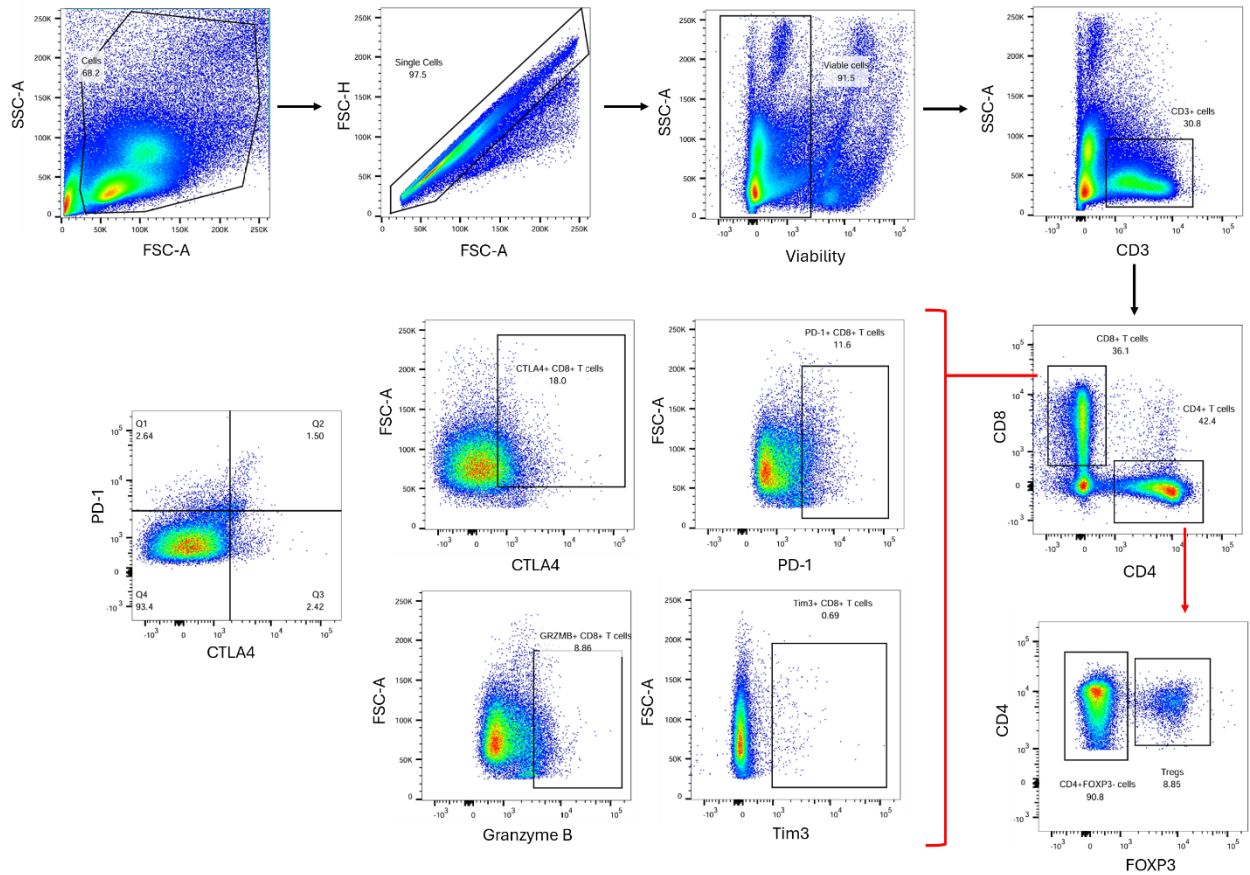

**Figure S3. Flow cytometry gating strategy for the murine lymph node panel.**

The gating strategy of the PD-1, CTLA4, Tim3, and granzyme B expression in the CD8+ T cell population was also applied identically to the regulatory T cells and CD4+ T cell population.
